# Success Criteria for Preclinical Testing of Cell-Instructive Hydrogels for Tendon Regeneration

**DOI:** 10.1101/2020.07.16.207274

**Authors:** Ryan C. Locke, Eden M. Ford, Karin G. Silbernagel, April M. Kloxin, Megan L. Killian

## Abstract

Tendon injuries are difficult to heal in part because intrinsic tendon healing, which is dominated by scar tissue formation, does not effectively regenerate the native structure and function of healthy tendon. Further, many current treatment strategies also fall short of producing regenerated tendon with the native properties of healthy tendon. There is increasing interest in the use of cell-instructive strategies to limit the intrinsic fibrotic response following injury and improve the regenerative capacity of tendon *in vivo*. We have established multi-functional, cell-instructive hydrogels for treating injured tendon that afford tunable control over the biomechanical, biochemical, and structural properties of the cell microenvironments. Specifically, we incorporated integrin-binding domains (RGDS) and assembled multi-functional collagen mimetic peptides (mfCMPs) that enable cell adhesion and elongation of stem cells within synthetic hydrogels of designed biomechanical properties and evaluated these materials using targeted success criteria developed for testing in mechanically-demanding environments like tendon healing. The *in vitro* and *in situ* success criteria were determined based on systematic reviews of the most commonly reported outcome measures of hydrogels for tendon repair and established standards for testing of biomaterials. We then showed, using validation experiments, that multi-functional and synthetic hydrogels meet these criteria. Specifically, these hydrogels have mechanical properties comparable to developing tendon; are non-cytotoxic both in 2D bolus exposure (hydrogel components) and 3D encapsulation (full hydrogel); are formed, retained, and visualized within tendon defects over time (two-weeks); and provide mechanical support to tendon defects at the time of injection and *in situ* formation. Ultimately, the *in vitro* and *in situ* success criteria evaluated in this study were designed for preclinical research to rigorously test the potential to achieve successful tendon repair prior to *in vivo* testing and indicate the promise of multi-functional and synthetic hydrogels for continued translation.

**IMPACT STATEMENT:** Tendon healing results in a weak scar that forms due to poor cell-mediated repair of the injured tissue. Treatments that tailor the instructions experienced by cells during healing afford opportunities to regenerate the healthy tendon. Engineered cell-instructive cues, including the biomechanical, biochemical, and structural properties of the cell microenvironment, within multi-functional synthetic hydrogels are promising therapeutic strategies for tissue regeneration. In this paper, the preclinical efficacy of multi-functional synthetic hydrogels for tendon repair is tested against rigorous *in vitro* and *in situ* success criteria. This study indicates the promise for continued preclinical translation of synthetic hydrogels for tissue regeneration.

## INTRODUCTION

Nearly half of the 26.3 million musculoskeletal injuries and procedures in the US each year involve tendons^1^. Although current treatment strategies for injured tendons (e.g., sutures and grafts) increase the acute mechanical function of tendon after injury, re-injury rates remain high as the challenge of tendon regeneration is not solely mechanical in nature^2–7^. Healthy tendon is not regenerated following injury and often heals via formation of fibrotic and mechanically weak scar tissue^7–9^. Therefore, new treatment strategies are needed to provide cellular instructions that shift the healing response from fibrotic to regenerative. Natural and synthetic hydrogels range in material type and functionality and have been used in *in vitro* and *in vivo* studies for tendon healing; yet, a broad range of outcome measures have been reported to determine their preclinical safety and success. A standardized list of *in vitro* and *in situ* success criteria to meet prior to *in vivo* testing would benefit researchers for the rigorous and systematic testing of hydrogels in mechanically-demanding environments like tendon repair.

Guidelines to evaluate biomaterials for tissue repair include those generally defined by the International Organization for Standardization (ISO 10993)^10^ and by the American Society for Testing and Materials (ASTM, e.g., ASTM F2900 and ASTM F2903). In the literature, expected outcomes have been reported related to tissue-engineered constructs for tendon regeneration *in vivo*^11–16^. However, success criteria for evaluating hydrogels designed for tendon repair prior to *in vivo* application are not clearly defined. A limitation of hydrogels compared to other tendon repair strategies is the ability of the hydrogel to be retained within the injury site long-term and also provide mechanical support while promoting tenogenic remodeling^17,18^. Integration of related metrics to evaluate material performance prior to *in vivo* implantation would facilitate design and translation of hydrogels for tendon repair.

Hydrogels, including collagen-^19^ and extracellular matrix (ECM)-based^20^ gels, are used as carriers to guide and deliver stem cells. However, the clinical efficacy and the role of transplanted cells in tendon repair remains ambiguous^21^. Promising new synthetic hydrogels may circumvent the need for stem cell delivery if these materials contain deliberately designed cell-instructive cues, such as cell integrin-binding domains, fibrous ECM-mimetic peptides, tunable mechanical properties, and matrix metalloprotease (MMP)-degradable linker peptides^17,22^. Engineered hydrogels, such as functionalized poly(ethylene glycol) (PEG) hydrogels, are designed for delivery of targeted cellular instructions via tunable control of mechanical and biochemical environments^23–26^. Such cell-instructive hydrogels and scaffolds are attractive alternatives to cell-based tendon therapies because these materials do not rely on allogenic cell use and therefore reduce cell-donor site morbidity and immune rejection^27–34^. Drawbacks of synthetic hydrogels, however, can include cytotoxicity (e.g., by-products of polymerization steps) or limited mechanical support compared to sutures and fibrous scaffolds^15,17,35^. Thus, developing innovative material designs that integrate bioactive structures may address these drawbacks and improve preclinical translation for tendon repair.

In this work, we developed and tested *in vitro* and *in situ* success criteria for hydrogels in mechanically-demanding environments specific for use in tendon repair. Criteria were determined via systematic review of the peer-reviewed literature. We performed validation experiments both *in vitro* and *in situ* using preclinical models of tendon injury. Here, photo-polymerized PEG-peptide thiol–ene hydrogels were used *i)* with well-defined homogenous nanostructure^25,36,37^ and *ii)* with fibrillar and collagen mimetic nanostructure. Through testing of synthetic hydrogels against the developed success criteria in the context of tendon repair, we established that multi-functional, cell-instructive synthetic hydrogels hold promise for continued translation for tendon repair.

## METHOD

### Multi-functional Collagen Mimetic Peptide (mfCMP) Hydrogel

Hydrogels were designed to be locally cell-degradable, accounting for the need for cell migration and cell-mediated ECM turnover in the context of *in vitro* and *in vivo* applications. Specifically, we incorporated into synthetic PEG-peptide hydrogels a matrix metalloproteinase (MMP)-degradable linker (degradable linker peptide), a variant of that found in collagen type-I known to degrade in response to a variety of MMPs including MMP-1 and MMP-2, which are upregulated during tendon repair^38,39^. As tendon ECM is primarily fibrous collagen^39,40^, the incorporation of collagen-like structure may improve tendon repair *in vivo*. We have therefore incorporated an assembled, multifunctional collagen mimetic peptide (mfCMP) into this hydrogel system to provide fibrous structure^23^. Here, to keep degradable content constant with and without mfCMP, a scrambled linker with reduced MMP-specific degradability (scrambled linker peptide) also was incorporated into the network and replaced with the mfCMP (a nondegradable peptide) when desired for increased structural content. Additionally, pendant RGDS (an integrin-binding peptide) and a fluorophore were incorporated into the hydrogel network to promote cell adhesion and enable imaging within tissues. The design of this material allows for independent control over modulus, structure, biochemical content, and degradability, which can individually be tuned for mimicking aspects of developing tendon, and enables *in situ* formation within tendon injuries.

### Success Criteria for Tendon Repair with Cell-Instructive Hydrogels

Most prior reports primarily focused on mechanical and biological outcomes to meet during *in vivo* tendon healing (Table S1)^11–16^. We developed *in vitro* and *in situ* success criteria by populating a list of the most common outcome measures performed on hydrogels intended for tendon repair (Supplemental Methods (SM): Systematic Reviews; Figures 1 and S1-4, Table S2). We then searched clinical literature of Achilles tendon and rotator cuff repair to identify the most common outcome measures used to define successful clinical repair (SM: Systematic Reviews; Figures 1 and S5, Table S3). We then cross-referenced these lists with ISO and ASTM standards (Figure 1 and Table S4 and Lists S1-3).

**Figure 1.**
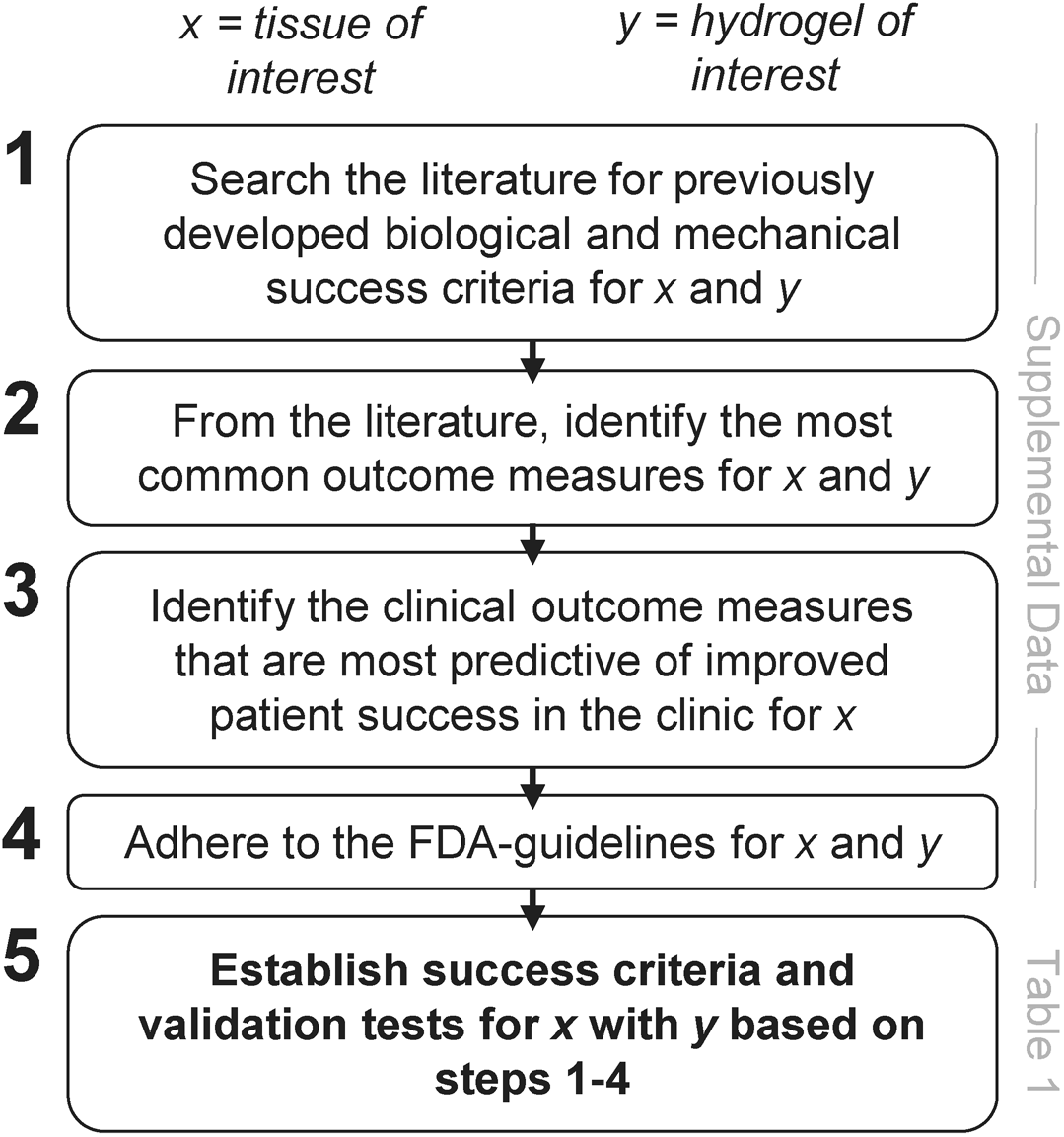
Study design and method. (1) Literature searches were performed for prior success criteria for tendon repair. Systematic reviews were performed to identify the most common outcome measures for (2) preclinical use of hydrogels for tendon repair and (3) clinical tendon repair. (4) FDA guidelines were cross-referenced. (5) Success criteria and validation tests were established based on the identified gaps in the literature and the findings from 1-4. Table 1 outlines the success criteria that we developed and tested in this study for tendon repair using hydrogels. See the supplemental data for outcomes from steps 1-4.

For the final criterion (Table 1), we performed validation experiments using the hydrogel system described above. The success criteria can be tailored toward the mechano-biological needs from tendon to another tissue (x) or hydrogel (y) using the 5-step method in Figure 1.

**Table 1.** *In vitro* and *in situ* success criteria to meet for the regeneration of injured tendons with in situ-formed, cell-instructive hydrogels.

|  | **Success Criteria** | **Tests\*** | **Description of this study** |
| --- | --- | --- | --- |
| **Material Properties** | In situ-formed with tunable & physiologically-relevant modulus | Hydrogel Mechanical Testing* | Material properties of developing tendon compared to gel (Figure 2) |
| **Degradation Profile** | Controlled material degradation profile | Enzyme Incubations* | Prior data validates that our materials allow controlled degradation. |
| **In Vitro Compatibility** | Maintain viability of both primary human and animal cells, while allowing molecular analyses | Individual Chemical Cytotoxicity | Culture cells with each hydrogel component to the media, and test viability (Figure 3A) |
|  |  | Bulk Cytotoxicity | Culture cells within 3-dimensional (3D) hydrogels and test viability (Figure 3B) |
|  |  | Morphology & Migration* | Culture cells within 3D hydrogels and perform imaging within the gel and IHC or RNA isolation (Figure 3B and S10) |
|  |  | Molecular Biology |  |
| **Material-Tissue Interface** | Visualize the interface or direct bond of the material to the tissue defect | Multi-scale Imaging*<br>Histology | Macroscale images of the hydrogel formed with a rat Achilles tendon defect (Figure 4) |
| **In Vivo Imaging** | Localize the material when applied in vivo | IVIS Tracking | IVIS image hydrogel with varying concentrations of AF647 (Figure 5) |
| **Long-Term Retention** | In situ-formed and retained within tissue defects long-term (weeks) | Physiologic Incubations<br>Fatigue Testing* | Create defect within rat Achilles tendon, form the hydrogel in the defect, and incubate the tendon-hydrogel construct under physiological conditions (Figure 6) |
| **Mechanical Support** | Provide mechanical support to tissue defects at the time of injection | Load-to-Failure Testing* | Create defect within rat infraspinatus tendon-bone attachment, form the hydrogel in the defect, and load-to-failure (Figure 7) |

\*This list does not include all possible tests, instead it includes the tests that were most commonly performed to validate the use of hydrogels for tendon repair. If there is a specific goal of the repair strategy – for example, to induce cell differentiation – then this list can be adapted to include differentiation markers (RNA/IHC) or cell morphology assays. Rheology is one of the most common mechanical tests for hydrogels. Fatigue and load-to-failure test protocols are context-dependent. Enzyme incubations, cell migration, multi-scale imaging (e.g., nano- to macro-scale), and fatigue testing were not performed in this study, however they are important for hydrogel success in tendon repair *in vivo* and are avenues for future studies.

## EXPERIMENT

### Experimental Design

#### Peptide and mfCMP Synthesis

Four peptides in total were synthesized on an automated peptide synthesizer with microwave capabilities (Liberty Blue; CEM Corporation, Matthews, NC), using solid phase peptide synthesis (SPPS) techniques according to a previously described protocol^23^. The two linker peptide variants used in this study included a degradable linker peptide (KK(**alloc**)G<u>GPQG↓IWGQ</u>GK(**alloc**)K) and a scrambled linker peptide (KK(**alloc**)GIQPGGWGQGK(**alloc**)K). Here, K(**alloc**) is an allyloxycarbonyl-protected lysine residue that presents a reactive handle for light-initiated thiol–ene click chemistry. In addition, a multifunctional collagen mimetic peptide (mfCMP, (PKG)_4_PK(alloc)G(POG)_6_(DOG)_4_, where O is a hydroxyproline residue) and an integrin-binding peptide (RGDS, K(**alloc**)GWG<u>RGDS</u>) were synthesized to impart fibrillar structure and cell adhesion, respectively (SM; Peptide Synthesis and mfCMP Synthesis). Collected fractions were frozen, lyophilized, and products were confirmed via electrospray ionization (ESI+) mass spectrometry (Xevo G2-S QTof; Waters Corporation, Milford, MA) (Figures S6-8). Stock solutions of all peptides were prepared in Dulbecco’s phosphate buffered saline (DPBS). mfCMP concentrations were determined by weight, and ultraviolet-visible (UV-VIS) spectrometry (NanoDrop™ 2000c Spectrophotometer; Thermo Fisher Scientific, Waltham, MA) was used to confirm stock solution concentrations of all other peptides via converting tryptophan absorbance at 280 nm to peptide concentration (Beer’s law, ε_280,Tryptophan_ = 5690 M^−1^ cm^−1^)^23^.

#### Hydrogel Design, Photopolymerization, and Rheometry

Four-arm PEG with hydroxyl end groups (PEG-4OH, JenKem, M_n_ ~ 20 kDa) was modified with thiol end groups as previously described^23,41^, producing hydrogel precursor PEG-4SH (SM; PEG-4SH Modification). PEG-4SH was diluted into DPBS (25mM thiol functional groups) and Alexa Fluor™ 647 C_2_ Maleimide (Thermo Fisher Scientific, Waltham, MA) was added targeting 0.5% thiol conjugation with the fluorophore (Figure 5A; 80μM AF647:16mM thiols in the final hydrogel precursor solution, 1-hour reaction, modified from manufacturer’s protocol). All hydrogels were fluorescently labeled with Alexa Fluor™ 647 (AF647) where 80μM AF647 was used for *in situ* imaging in tendon, unless otherwise stated.

Hydrogel precursor solutions were prepared prior to photopolymerization and contained 8wt% PEG-4SH, degradable linker peptide, scrambled linker peptide (1.25mM, 2.5mM alloc; 0mM mfCMP condition) or mfCMP (2.5mM, 2.5mM alloc; 2.5mM mfCMP condition), and RGDS such that thiol and alloc functional groups were stoichiometrically balanced (16mM thiol:16mM alloc) unless otherwise stated. The RGDS peptide was incorporated at 2mM (2mM alloc), and degradable content was kept constant with 5.75mM degradable linker peptide (11.5mM alloc) incorporation. All solutions included 2.2mM LAP photoinitiator synthesized by previously published protocols (SM; Lap Synthesis)^42^. Hydrogel precursor solutions (20μl) were transferred to molds (sterile 1mL flat-plunger syringes with tips removed; Thermo Fisher Scientific, Waltham, MA) and irradiated with cytocompatible doses of collimated long wavelength UV light for 4-minutes (365nm, 10mW/cm^2^; Exfo Omnicure Series 2000 light source, 365nm bandpass filter, Excelitas Technologies Corp., Waltham, MA)^23^.

Hydrogels (8wt%; 80μM AF647; 0mM and 2.5mM mfCMP conditions) were formed *in situ* on a AR-G2 rheometer with a quartz plate UV-curing light accessory (TA Instruments, New Castle, DE). A liquid-filled light guide was used to attach the UV curing light accessory to a UV-visible light source with bandpass filter (Exfo Omnicure Series 2000 light source, 365nm bandpass filter, Excelitas Technologies Corp., Waltham, MA). Hydrogel precursor solutions (10μL) were pipetted onto the quartz plate, and an 8mm flat plate top was lowered to a 160μm gap height. Storage modulus (G′) measurements were collected at room temperature with 1% strain and 2rad/s frequency (within the linear viscoelastic regime). Measurements were taken for 60-seconds prior to irradiation followed by 4-minutes of irradiation (365nm, 10mW/cm^2^) to monitor complete hydrogel formation, at which point the final modulus measurements were determined. G′ was converted to elastic modulus (Young’s modulus, *E*) using rubber elasticity theory^43^ assuming a Poisson’s ratio of 0.5 for these elastic, incompressible hydrogels and compared to the elastic modulus of developing chick tendon^44^ (Figure S9, SM; Modulus Conversion). Note that two minutes of irradiation was determined to be sufficient for complete gelation and was used in *Individual Chemical Cytotoxicity* and *Mechanical Testing* experiments.

#### Cell Culture Experiments

One, adult female Long Evans rat was euthanized via carbon dioxide asphyxiation. Immediately, long bones were dissected and bone marrow mesenchymal stem/stromal cells (rBMSCs) isolated using a modified version of a published protocol (SM; rBMSC Culture)^45^. rBMSCs were cultured on a 100mm^2^ petri dish in growth media (Dulbecco’s modified eagle medium (DMEM), and 1% penicillin, streptomycin, amphotericin, Genesee Scientific; and 10% fetal bovine serum (FBS), Atlanta Biologics) at 37°C and 5% CO_2_. Media was replaced every 2 days. rBMSCs were passaged at 1wk (Passage 1, P1) and cultured until 85% confluency before cytotoxicity studies described below (performed at P2).

Human mesenchymal stem/stromal cells (hMSCs; Lonza, Basel, Switzerland) derived from bone marrow were expanded on tissue culture treated polystyrene flasks (182cm^2^; CELLTREAT, Pepperell, MA) at 37°C and 5% CO_2_ using standard sterile technique. hMSCs were thawed and cultured in growth media (low glucose DMEM with penicillin, streptomycin, Fungizone, and 10%v/v FBS, all from Thermo Fisher Scientific, Waltham, MA; with 1ng/mL bFGF; PeproTech, Rocky Hill, NJ). Media was replaced every 3-4 days, and hMSCs were passaged once cells reached 85% confluency. All hMSC experiments were conducted with cells at P6-7.

For experiments, hMSCs and rBMSCs were plated at 15,000cells/cm^2^ in 96-well plates and cultured overnight prior to treatment. Cells were exposed to one of five treatments: 1) negative control (2-minute PBS incubation), 2) long wavelength UV light (2-minutes in PBS; 365nm; 10mW/cm^2^), 3) blue light (2-minutes in PBS; 400-500nm bandpass filter; 10mW/cm^2^), 4) radical exposure (2-minutes in PBS with 0.2mM LAP; 365nm; 10mW/cm^2^), and 5) all hydrogel components (2-minutes in PBS with 0.2mM LAP, 0.5mM degradable linker peptide and 1mM L-Cysteine, 365nm, 10mW/cm^2^) (Exfo Omnicure Series 2000 light source with liquid light guide with collimating lens and respective bandpass filter, Excelitas Technologies Corp., Waltham, MA). L-Cysteine was used in place of PEG-4SH to mimic hydrogel formation conditions without gelation, allowing cell exposure to the photoinitiated thiol–ene reaction. After treatment, cells were allowed to recover overnight in growth media. alamarBlue™ (Thermo Fisher Scientific, Waltham, MA) was used to measure metabolic activity (SM; alamarBlue™) and the fluorescence was measured on a plate reader (ex/em: 570/585; Synergy 2 plate reader; BioTek, Winooski, VT).

#### 3D Cell Encapsulation, Viability, and Collagen Deposition

hMSCs were encapsulated within hydrogels that were not fluorescently labeled (6wt%; 0μM AF647; 0mM and 2.5mM mfCMP conditions; SM; 3D Cell Encapsulation)^23^. Two hydrogel conditions were prepared: without mfCMP (2mM RGDS, 5mM degradable linker peptide) and with mfCMP (2mM RGDS, 3.75mM degradable linker peptide, 2.5mM mfCMP). hMSCs were suspended in the hydrogel precursor solution at 3.75 × 10^6^cells/mL, and 20μL cell-encapsulated hydrogels were formed in syringe molds following procedures described above, under sterile conditions. Hydrogels were transferred to a non-treated 48-well plate with bFGF-free media (500μL; SM; 3D Cell Encapsulation) and incubated at 37°C and 5% CO_2_ using standard sterile technique. Media was replaced every 2 days.

Encapsulated hMSC viability was assessed 3-days after encapsulation using a commercially available kit (Live/Dead™ mammalian cell viability kit, Thermo Fisher Scientific, Waltham, MA). Z-stacks (200μm) were acquired using a confocal microscope (Zeiss LSM 800; Zeiss, Oberkochen, Germany), and orthogonal projections were used to count the number of live and dead cells in ImageJ^46^.

After 16-days of culture, hydrogels were fixed in 4% methanol-free formaldehyde (PFA; Thermo Fisher Scientific, Waltham, MA) and washed in bovine serum albumin (BSA, MilliporeSigma, Burlington, MA). Hydrogels were incubated in a BSA blocking solution then washed in 0.2%v/v TWEEN®20 (MilliporeSigma, Burlington, MA). Triton™ X-100 (Thermo Fisher Scientific, Waltham, MA) was used to permeabilize cells for 30-minutes. Hydrogels were then incubated sequentially with rabbit-anti-human collagen type-I antibody (Abcam, Cambridge, UK), fluorescently-labeled goat-anti-rabbit secondary antibody (Thermo Fisher Scientific, Waltham, MA), and Hoechst (Thermo Fisher Scientific, Waltham, MA). Z-stacks (200μm) were acquired via confocal microscope and orthogonal projections were produced for analysis. Toward studying behavior of cells in response to these materials, we also developed a protocol for extracting RNA from hydrogels with low numbers of cells (e.g. hMSCs, SM; RNA Isolation from Synthetic Hydrogels; Figure S10).

#### Ex Vivo studies

Achilles tendons of adult, female Long Evans rats that were euthanized and fresh frozen were used in this study (fresh frozen, N=8). A 1mm biopsy punch was used to make a circular defect in the center third of the Achilles tendon midsubstance ~3mm above the Achilles-calcaneus attachment (n=6, defect group). For complete laceration, a #11 scalpel blade was used ~3mm above the Achilles-calcaneus attachment to make a complete laceration (n=6, laceration group). Two, non-damaged tendons were used as uninjured controls (n=2).

Following laceration, the two tendon stumps were approximated end-to-end before hydrogel formation (Video S1). Hydrogel precursor solutions (8wt%; 80μM AF647; 0mM and 2.5mM mfCMP compositions) were pipetted into the tendon defects and photopolymerized with a collimated lamp following procedures described above (365nm, 10mW/cm^2^, SM; Formation in Achilles Defects). The interface between the tendon and hydrogel was imaged with a digital camera (Video S1; TG-860, Olympus). Tendons were stored in conical tubes with PBS and penicillin, streptomycin, and amphotericin.

To probe what fluorophore concentration was sufficient for *in situ* imaging, AF647-labeled hydrogels (20μl) were formed with 10μM, 50μM, 80μM, or 100μM AF647; immediately swelled in PBS for 2-hours; and then imaged. Hydrogels were imaged using an IVIS (Perkin Elmer) with or without rat skin between the hydrogel and camera.

Tendon-hydrogel constructs (n=12 tendons) were agitated on an oscillating rocker (1Hz, 22°C unless otherwise noted) for two weeks to assess long-term retention of hydrogels. Tendon-hydrogel constructs were imaged using the IVIS. Images were taken using the IVIS after 2-, 4-, and 6-hours, and 1-, 3-, 5-, 7-, and 14-days of oscillation. The radiant efficacy and integrity (defined as present or not present; broken or not broken) were recorded.

#### Mechanical Testing of Hydrogels within Partial-Thickness Rotator Cuff Injury Model

The rotator cuff tendons of fresh-frozen adult, female Long Evans rats were used in this study (N=6 rats with defects, N=1 intact control). Partial-thickness defects were made bilaterally at the center of the infraspinatus tendon attachment using a 300μm biopsy punch as previously published (n=12)^47,48^ and humeri were potted in poly-methyl methacrylate (Ortho-Jet BCS, Lang Dental, Wheeling, IL)^48^. Defects received either hydrogel (n=6) or saline (empty control group, n=6) for paired comparisons. Hydrogels without mfCMP (10μl; 10wt%; 2 mM RGDS; 9mM degradable linker peptide) were formed by photopolymerization (2-minutes, 365nm, 10mW/cm^2^). After hydrogel formation, all samples from both groups were incubated in PBS at 37°C for 1-hour to simulate physiological conditions and induce hydrogel swelling. Samples were subjected to uniaxial tensile tests as follows: 0.2N tare load applied with 5 preconditioning cycles (±0.05mm) and load to failure at 0.01mm/s. The cross-sectional area (assumed circular) and gauge length were measured at the start of each test and mechanical properties were calculated using a custom MATLAB script^48^.

#### Statistics

Storage moduli and 3D culture viability were compared between 0mM CMP and 2.5mM CMP hydrogels using unpaired Student’s t-tests. Metabolic activity of all the treated groups was compared using one-way ANOVA with Tukey’s post-hoc test. Radiant efficiency was compared between the four individual groups using repeated measures (time) two-way ANOVAs with Tukey’s post-hoc test. Because no differences were observed between groups, all groups were then combined and analyzed by repeated measures (time) one-way ANOVA with Tukey’s post-hoc test, and the grouped data were fit using a one phase decay. Metabolic activity (background:0, control average:100) and radiant efficiency (detectable limit:0, 2-hour average:100) were linearly normalized for graphical representation. Mechanical properties of the injured infraspinatus attachment with and without hydrogels were compared using paired Student’s t-tests.

### Experimental Results

#### Success Criteria

We developed a list of *in vitro* and *in situ* success criteria for evaluation of hydrogels in mechanically-demanding environments prior to their use within *in vivo* applications (Figure 1 and Table 1) and tested each of these criterion with synthetic, multi-functional hydrogels. We identified 12 outcome measures that were most frequently reported in the literature, and, of these, all were also reported in standard guides (Tables S1-4). Preclinical outcomes concentrated on histological, biomechanical, and biochemical assays, while clinical outcomes focused on repair integrity through post-operative imaging, joint function, and surgical complications and were similar between tendons (Table S2-3).

#### Hydrogel Material Properties

Crosslinked hydrogels with and without assembled mfCMP (Figure 2A) reached full gelation after two-minutes of irradiation with low-intensity, long wavelength UV light (≤1% change in modulus) (Figure 2B). We observed no significant difference in *in situ* elastic modulus between hydrogels with mfCMP (10.5±0.7 kPa) compared to those without mfCMP (10.7±0.3 kPa), and both were in the range of published values for developing chick tendon^44^ (Figure 2B). After equilibrium swelling, owing to water uptake, the elastic modulus of the hydrogels decreased to 4.4±0.3 kPa (SM; Equilibrium Swollen Gel Modulus Measurements, Figure S9), which is still within the range of developing chick tendon^44^.

**Figure 2.**
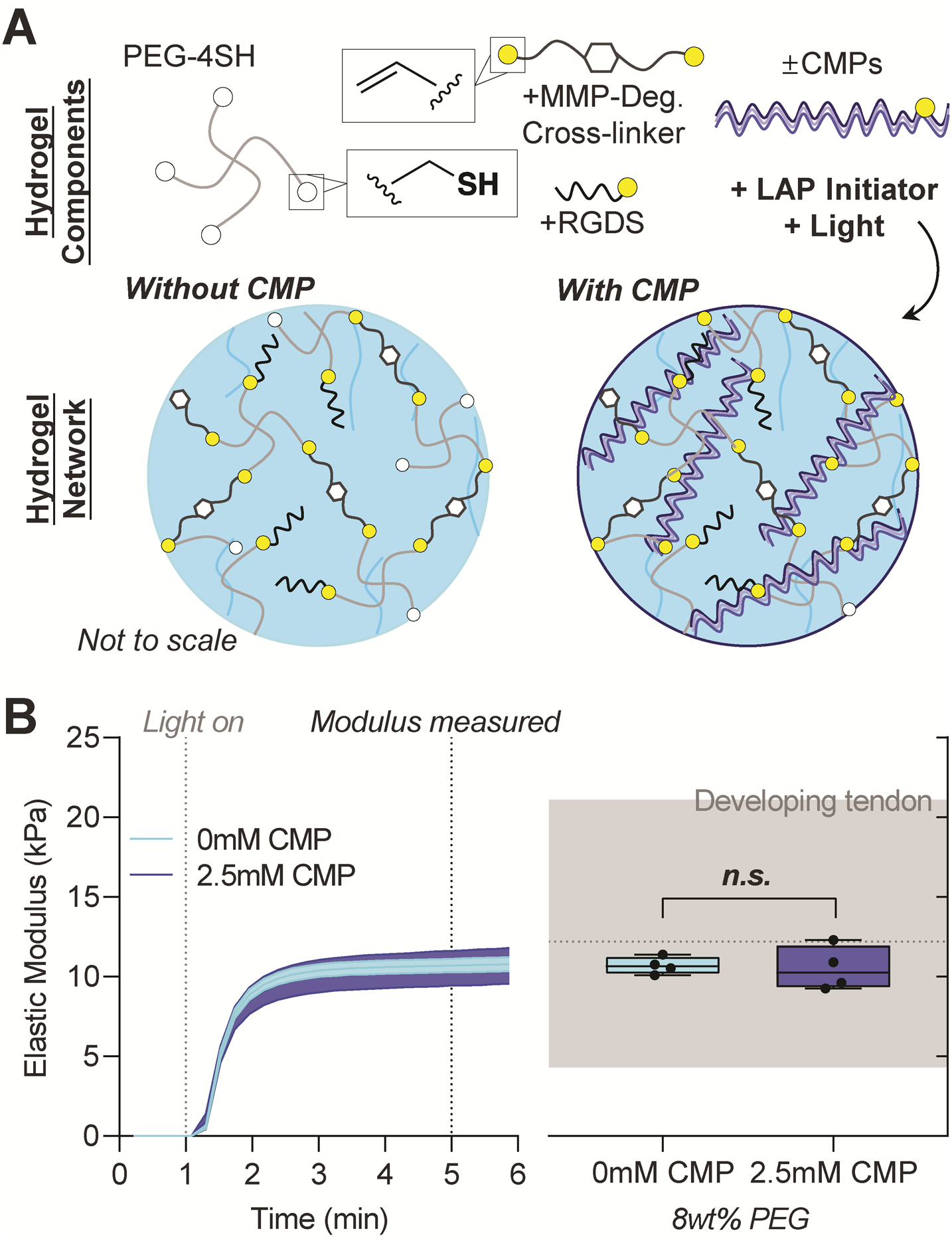
Hydrogel design, photopolymerization, and mechanical properties. (A) Hydrogel precursors, including PEG-4SH, a MMP-degradable linker peptide, RGDS cell-adhesive peptide, and assembled fibrillar mfCMPs, were combined with a photoinitiator (LAP) and then irradiated by long wavelength UV light (10 mW/cm^2^ at 365 nm) to form hydrogel networks. (B) Storage modulus was monitored during *in situ* formation and converted to elastic modulus using rubber elasticity theory for ease of comparison to the mechanical properties of tissues (left, mean ± standard deviation). Elastic moduli of hydrogels formed *in situ* with and without mfCMP were measured after 4-minutes of irradiation and are similar to the elastic modulus of developing chick tendon (right, gray bar; 4.3-21.1kPa, dotted line: 12.2kPa at embryonic day HH41). Data are presented as median ± 95% confidence interval^44^.

#### Cell Compatibility

In 2D culture (Figure 3A), doses of light required for hydrogel formation (either blue and long wavelength UV light) with or without all individual components for hydrogel formation did not influence cell metabolic activity (Figure 3A). However, without monomers to scavenge radicals, irradiation in presence of photoinitiator alone led to decreased metabolic activity and differences in cell morphology and number (Figure 3A). In 3D cell encapsulation (Figure 3B), the viability of hMSCs was ~80% both with and without mfCMP after 3 days (Figure 3B), further supporting cytocompatibility of hydrogel formation. After long-term encapsulation, collagen type-I localization was observed in the pericellular matrix and hMSCs were more elongated in the presence of mfCMP (Figure 3B). Further, RNA yield was acceptable from individual hydrogels at 3- and 19-days following hMSC 3D culture (Figure S10).

**Figure 3.**
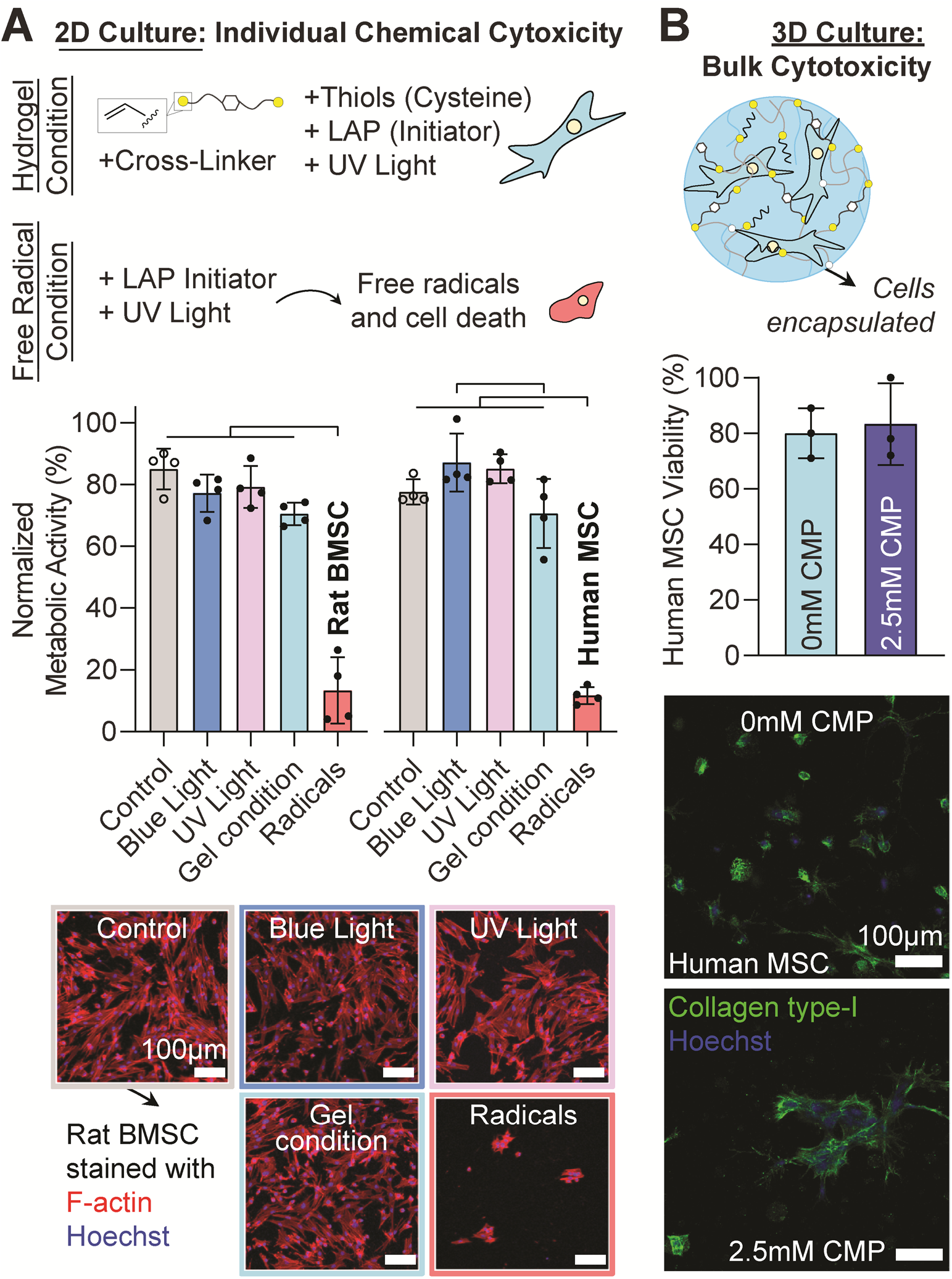
Hydrogel conditions are compatible with human and rat cells in 2D and 3D culture. (A) 2D cell culture schematic of the hydrogel and free radical culture conditions. Normalized metabolic activity of rat and human MSCs in 2D culture after exposure to different aspects of hydrogel formation, and example monolayer images of rat MSCs stained for F-actin (red) and nuclei (Hoechst, blue). (B) 3D cell encapsulation schematic, hMSC viability after 3-days of 3D encapsulation, and human collagen type-I immunostaining of hMSCs within hydrogels after 10-days of 3D encapsulation (6wt% gel). Data are presented as mean ± standard deviation. Black bars: significant difference between groups (p<0.05).

#### Tissue-Material Interface

Hydrogels were formed at the defect site of full-thickness defects and complete lacerations through rat Achilles tendons (Figure 4). The hydrogel prevented separation between the tendon stumps following complete laceration (Videos S1). The concentration of maleimide-functionalized AF647 used within the hydrogel precursor solution was detectable with a concentration of AF647 at 80μM underneath rat skin by imaging with IVIS, whereas less than 50μM was not detectable (Figure 5B).

**Figure 4.**
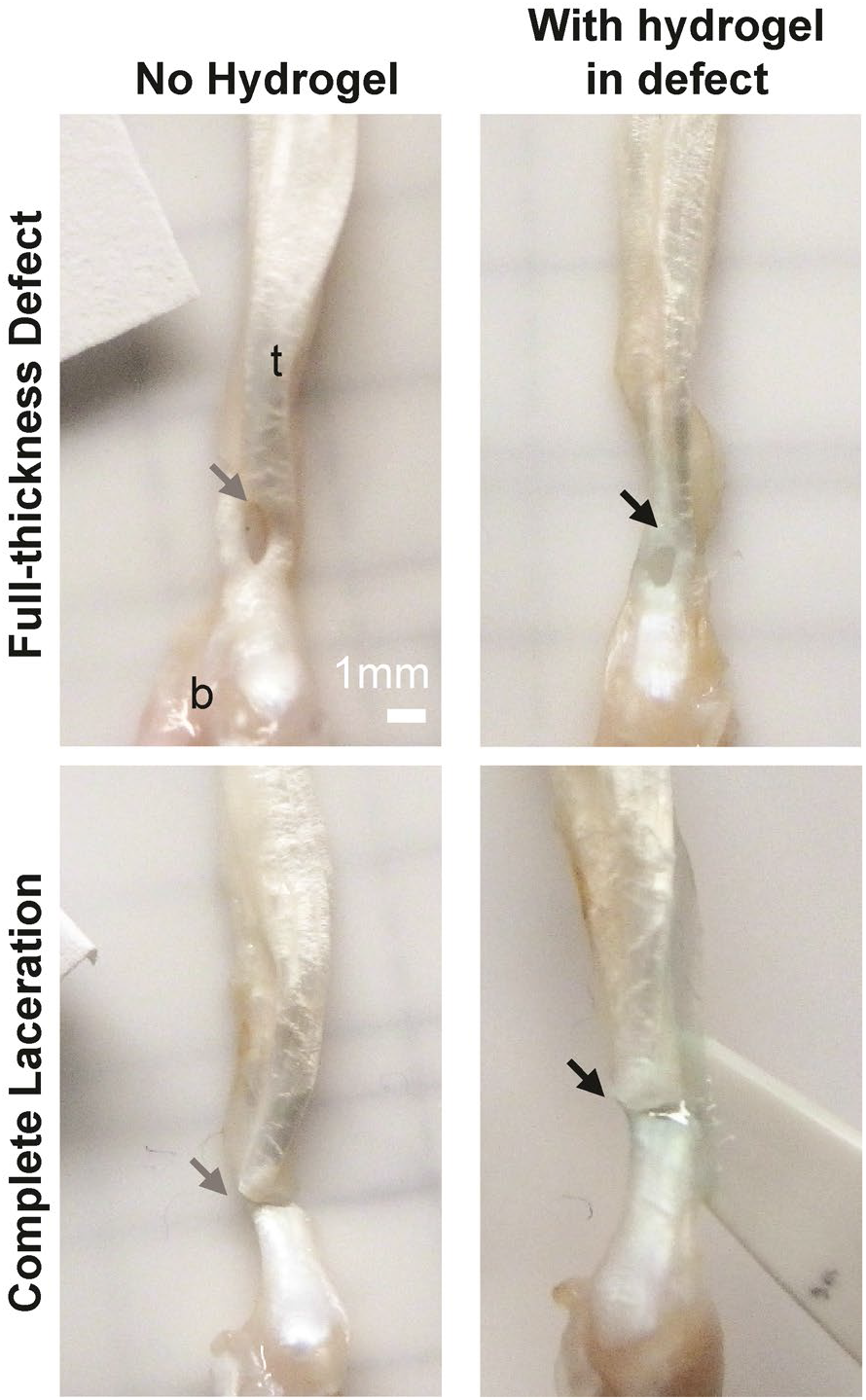
Macroscopically, a physical interface between tendon defects and hydrogel formed after photopolymerization. To visualize the tendon-hydrogel interface, full-thickness and complete laceration defect models in the rat Achilles tendon were imaged before (no hydrogel, gray arrows) and after hydrogel photopolymerization within the defect sites (with hydrogel in defect, black arrows). t: tendon, b: bone (calcaneus). See the supplemental video for further visualization. Scale bar: 1mm.

**Figure 5.**
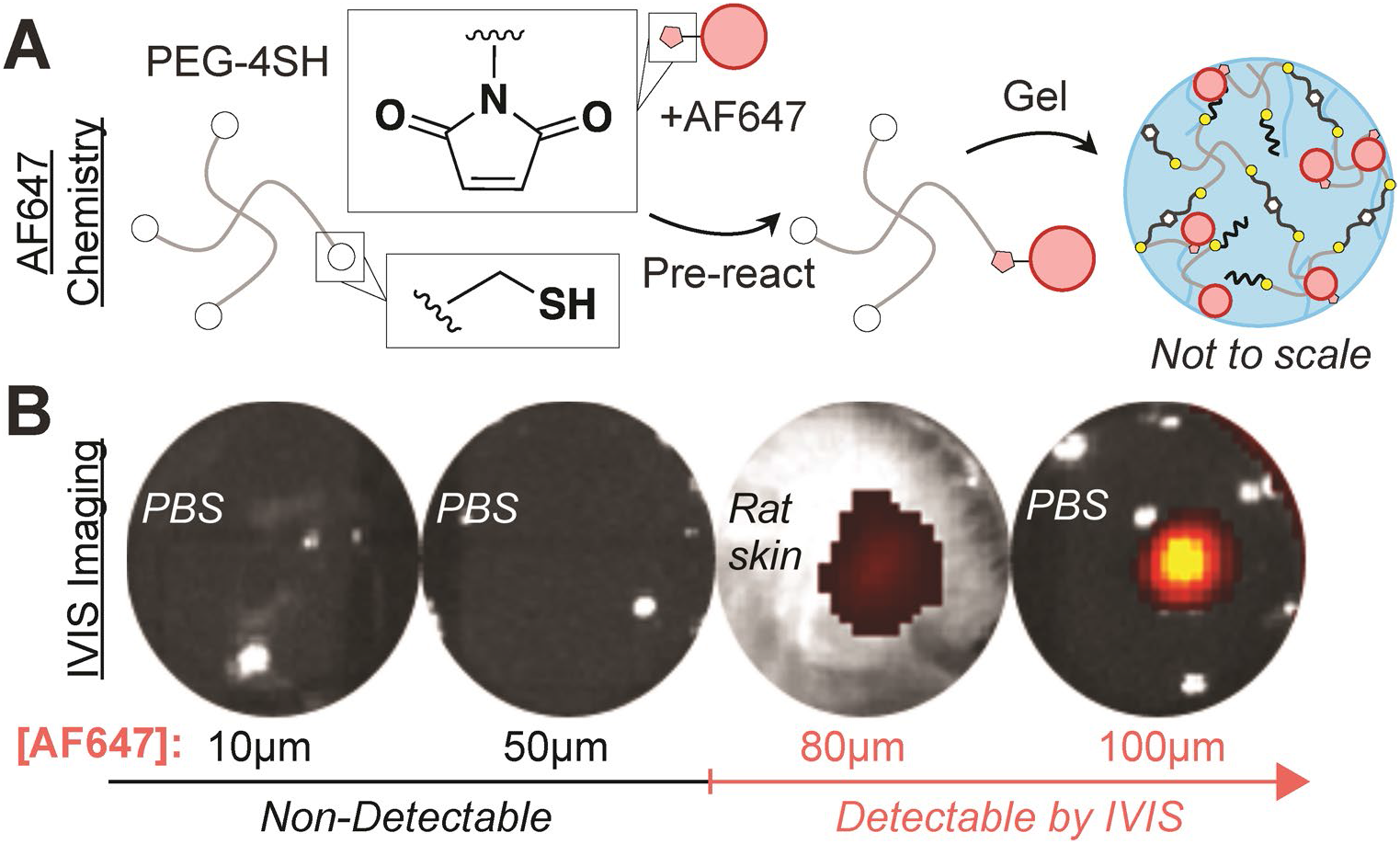
*In situ* imaging of hydrogels using IVIS. (A) AF647 was pre-reacted with PEG-4SH prior to photopolymerization to form fluorescent hydrogels. (B) IVIS images of fluorescent hydrogels with titrated concentrations of AF647.

#### Long-Term Retention

In Achilles full-thickness defects (Figure 6A), hydrogels with and without mfCMP remained in place and intact for over 2-weeks with fluid agitation (Figure 6B). For the complete laceration group, all tendon-hydrogel constructs remained intact at 3-days (Figure 6B). However, the majority of the hydrogels broke by 14-days at the tendon-hydrogel interface (Figure 6B) and 33% (n=2) remained fully intact (not shown). Notably, for all constructs that broke, the hydrogel remained adhered to the tendon stumps (Figure 6B). The radiant efficiency of each hydrogel was quantified and used as a readout of fluorophore loss due to potential degradation over time from agitation. No differences in AF647 radiant efficiency were detected between either of the hydrogel or defect groups, suggesting similar fluorophore loss and degradation rates with or without mfCMP (Figure 6C). With all hydrogels grouped together, the half-life of AF647 was nearly 5-days and the radiant efficiency remained well above the detectable limit after 2-weeks of incubation, indicating successful long-term retention and minimal degradation under agitation (Figure 6C).

**Figure 6.**
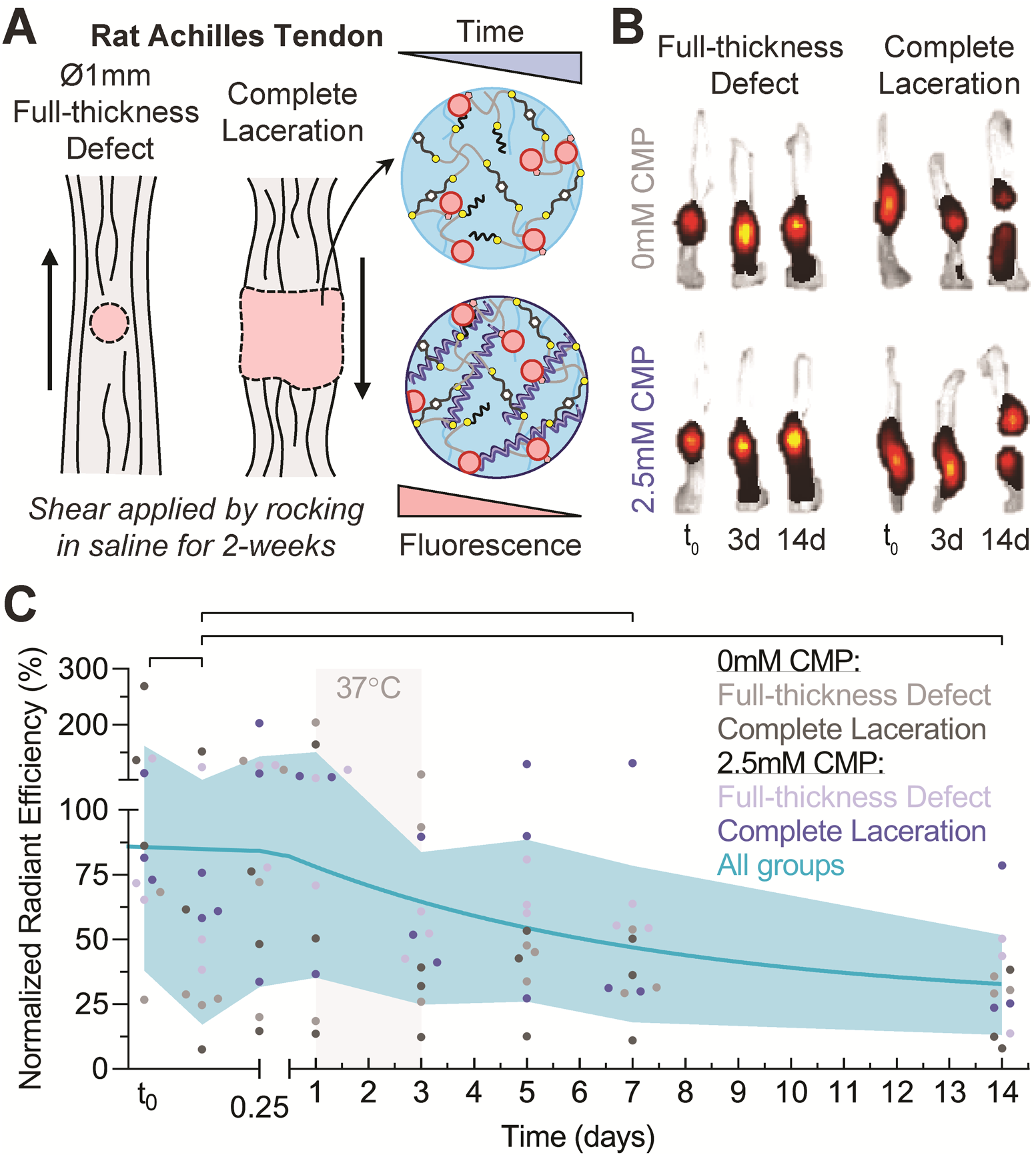
Hydrogels remained adhered to tendon defects for long-term under fluid agitation. (A) Schematic of Achilles tendon defect models and hydrogel rocking experiments. (B) Representative heat maps of AF647-hydrogel signal intensity for full-thickness defects and complete laceration tendon-hydrogel constructs with or without mfCMP after 2hrs, 3-days, and 14-days of rocking in PBS. (C) Quantified signal intensity (normalized radiant efficiency) of all hydrogels as individual groups (dots) and grouped together (shaded blue, standard deviation of all groups). The half-life of AF647 signal within hydrogels was approximately 5-days (blue line, one phase-decay fit). Samples were incubated at 37°C from 1-day to 3-day time points to determine the effect of physiological temperature (gray bar). Black bars: significant difference between time points for grouped data (p<0.05).

#### Mechanical Support

Inclusion of hydrogels in a rat model of partial-thickness rotator cuff attachment injury^47,48^ resulted to increased ultimate load in tension (24.3±4.1N) compared to empty defect controls (21.1±3.6N) but did not fully recover the mechanical properties of intact attachments (Figure 7A-B). No other mechanical properties measured were significantly different between groups (Figure S11). One sample was excluded and not tested due to scalpel damage during dissection.

**Figure 7.**
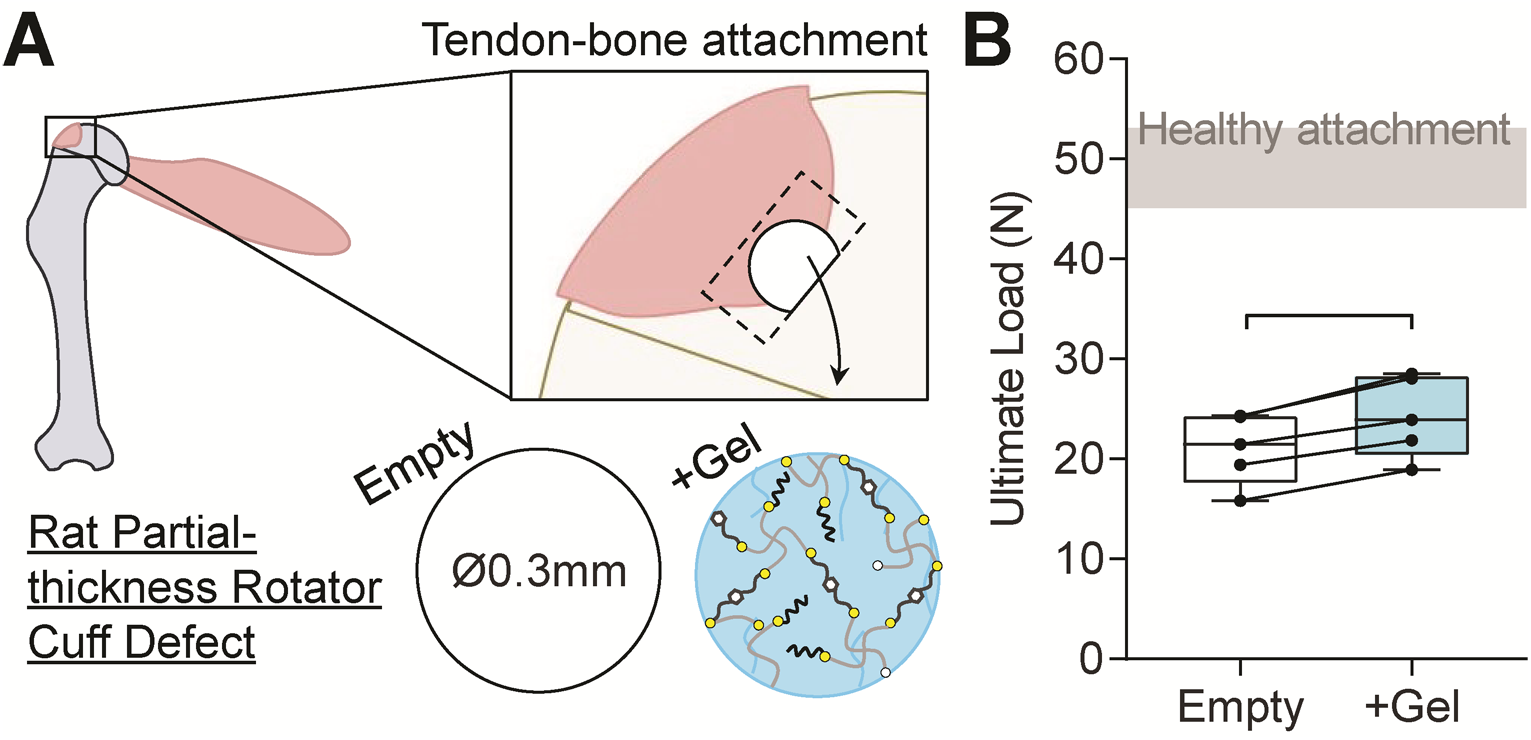
Hydrogels provide mechanical support upon *in situ* formation in defects. (A) Schematic of the rat partial-thickness rotator cuff defect model and injection site of the hydrogel precursor solution for *in situ* photopolymerization within the defect. (B) Ultimate load of empty and hydrogel-filled defects with respect to non-injured tendon (intact healthy attachment) and paired-limb comparison between empty and hydrogel groups (lines between individual data points). Data are presented as median ± 95% confidence intervals. Black bars: significant difference between groups (p<0.05).

## DISCUSSION

In this study, we first developed success criteria and then experimentally demonstrated that PEG-based hydrogels meet many of these criteria. Synthetic hydrogels offer opportunities for deliberate and tunable control over the bio-physical and -chemical cues for directing cell function and fate during healing. Specifically, the material stiffness, structure, and degradability are known to direct the behavior of stem cells, macrophages, and tendon fibroblasts^49–53^. Additionally, stiffness and degradability do not only direct cell differentiation^54,55^, but also may influence invasion^32^. Biochemical cues such as receptor-binding peptides and CMPs have been shown to modulate cell bioactivity^56,57^. For example, synthetic PEG-peptide-based ECM-mimics that contain integrin-binding CMPs have been shown to direct MSC differentiation toward tenogenic, chondrogenic, and osteogenic lineages^58,59^, and tenogenic-differentiation of MSCs may improve *in vivo* tendon repair^60^. Our established success criteria and a set of experimental approaches for preclinical studies rigorously assessed the efficacy of hydrogels using a combination of *in vitro*, *ex vivo*, and *in vivo* applications.

Beyond the provisional synthetic matrix of hydrogels, ECM secreted by cells, such as collagen type-I, is a tunable factor within engineered hydrogels and can influence hydrogel success^61–63^. In this work, we observed collagen type-I in the pericellular matrix of hMSCs in 3D culture with or without mfCMPs yet with significantly increased cell elongation in the presence of assembled mfCMPs, representing a cellular response to mfCMPs that may be a function of both ECM composition and structure. Bio-orthogonal labeling of hydrogels may be useful to determine the spatiotemporal effects of the cell-secreted ECM in hydrogels^62^ *in vitro* and in preclinical models *in vivo* ^64,65^.

Collagen-based hydrogels for the delivery of biologic supplements has been shown to support healing of the torn human anterior cruciate ligament^66,67^, lending substantial clinical evidence that hydrogels are retained in and suitable for healing of mechanically-demanding environments. While a combination of suture repair with hydrogel delivery may be necessary to maintain mechanical support^66^, we demonstrated that PEG-peptide hydrogels without suture repair can provide mechanical support to lacerated tendons at the time of injection. Further, these hydrogels, which we showed have similar mechanical properties to ECM of developing tendon^23,44^, offer tunable control over biochemical cues and enable a more targeted approach to improve healing. Indeed, in preclinical studies in the literature to date, hydrogels that incorporate mechano-biological cues of tendon development^58,68^, healthy tendon ECM^69–71^, or the ECM involved in tendon regeneration^72^ have shown benefits for tendon healing both *in vitro* and *in vivo.* Future research defining the ECM present in healthy and regenerated tendons, as well as further insights into the mechanical cues present during tendon development^44,73^ and post-natal growth^74^, may help to clarify the mechanisms of action for improved healing using these hydrogels.

We suggest that multiple, individual experiments for each criterion with carefully selected controls be performed to test materials against these success criteria. The success criteria and validation experiments should be tailored for the needs of the material prior to experimental testing. Rigorous evaluation criteria of hydrogels for tendon repair are essential for reducing rates of re-injury. Incorporating the success criteria we have defined here, along with adherence to other validated standards, into basic science investigations of hydrogels for tendon repair may increase the likelihood for successful repair and reduce the time from bench-to-bedside.

## ACKNOWLEDGEMENTS

This research was supported by Eunice Kennedy Shriver National Institute Of Child Health & Human Development of the National Institutes of Health (NIH) (K12HD073945), University of Delaware Research Foundation Strategic Initiatives (19A00297), and Delaware Clinical and Translational Research ACCEL (DE-CTR ACCEL) grant funded by an Institutional Development Award (IDeA) from the National Institute of General Medical Sciences (NIGMS) of the NIH (U54GM104941). E.M.F. was supported in part by a fellowship from the National Science Foundation (NSF) SBE2 IGERT Program (1144726). Instrumentation support was provided in part by the Delaware COBRE programs supported by grants from NIGMS of the NIH (P20GM104316, 5 P30 GM110758-02), University of Delaware NMR and Mass Spectrometry Core facilities, and the groups of Wilfred Chen and Dawn Elliott.

## DECLARATIONS

None of the authors of this manuscript have any competing interests to declare; therefore, no competing financial interests exist for any of the authors.

## Notes

### Competing Interest Statement

The authors have declared no competing interest.

