## Supplemental Data for "Success Criteria for Preclinical Testing of Cell-Instructive Hydrogels for Tendon Regeneration"

Tissue Engineering, Part C, Methods 2020

*Supplemental: Tables 1-4, Lists 1-3, Methods, Figures, and References*

[1]University of Delaware
Department of Biomedical Engineering
Newark, Delaware, USA

[2]University of Delaware
Department of Chemical & Biomolecular Engineering
Newark, Delaware, USA

[3]University of Delaware
Department of Physical Therapy
Newark, Delaware, USA

[4]University of Delaware
Department of Materials Science & Engineering
Newark, Delaware, USA

[5]University of Michigan Medical School
Department of Orthopaedic Surgery
Ann Arbor, Michigan, USA

[*]Shared First Authors
[#]Corresponding Authors

### Supplemental Tables

**Supplemental Table 1.** Previously reported outcome measures for determining preclinical safety and success for tendon repair using tissue-engineered approaches.

| Previous *In Vivo* Success Criteria | Reference |
|---|---|
| Functional Tissue Engineering: Biomechanical Criteria<br>• Measure *in vivo* applied forces (e.g., gait analysis)<br>• Aim to "best match" the sub-failure and failure properties of healthy tissue<br>• Select and prioritize mechanical outcomes<br>• Set quality standards for 'how good is good enough?'<br>• *In vivo* integration of material and native tissue/cells<br>• Pre-implantation, mechanical stimulation using bioreactors can improve structural and mechanical properties of tissue-engineered tendons<br>• Computational modeling of tissue healing can inform selection of time points for reducing animal numbers | [1–3] |
| Developmental Biology Perspective<br>• Target molecular markers of tendon development and determine their efficacy as therapeutics for tendon healing | [4] |
| Development Towards Successful Clinical Products<br>• Biodegradability: Degradation rate should be tuned to promote tissue healing and function; degrading too quickly may lead to poor structural support and too slowly may lead to stress-shielding or poor cell invasion.<br>• Biocompatibility: Immune response should not result in host rejection but may be tuned to promote integration into host tissue<br>• Processability: Match tissue size and shape with architectural structure of defect to mimic native tissue<br>• Mechanical strength: Provides mechanical integrity with applied loading and can maintain range of motion and frictionless movement/gliding<br>• Biofunctionality: Stimulate regeneration; mechanically stable; ability to be vascularized and allow nutrient diffusion; assist cell migration, proliferation, and differentiation until complete healing; avoid adhesion formation; assist supplement deliver<br>• Others: Material should be sterilizable, shelf-stable for long-term storage, and easy to handle in the clinic/operating room | [5] |
| Functional Tissue Engineering: Biological Criteria<br>• Material properties should be able to withstand estimated *in vivo* forces and stiffness<br>• If used in conjunction with stem cells, indicators of tenogenic differentiation (e.g., presence of Scx+ cells) should be observable<br>• Restored alignment and hierarchical ultra-structure of tendon and tendon-bone attachment | [6] |

**Supplemental Table 2.** Systematic review of the most common *in vitro* and *in vivo* preclinical outcome measures of hydrogels for tendon repair.

| Category | Top Outcomes | Percentage of n=87[7–93] |
|---|---|---|
| **Hydrogels** | | |
| Natural (N) | Collagen | 23% (53% total N) |
| Synthetic (S) | PEG | 13% (15% total S) |
| Natural and synthetic | N/A | (32% total dual N & S) |
| Gel formation | Self-assembly | 52% |
| | Light & chemical | 17% tie |
| Fiber-reinforced | No fiber | 77% |
| | Micro-fiber | 14% |
| | Nano-fiber | 9% |
| **Both *in vitro* and *in vivo*** | | **33% (100% total)** |
| ***In vitro* only** | | **48% (82% total)** |
| Outcome measures | Viability | 60% |
| | Bio-molecular & -chemical | 44% tie |
| | Proliferation or migration | 40% |
| | Cell morphology | 39% |
| | Electron microscopy | 33% |
| | Differentiation | 30% |
| | Degradation or release | 26% |
| Cell type | Stem cell & fibroblasts | 34% |
| | Fibroblasts | 29% |
| | Multiple cell types | 13% |
| | Other | 6% |
| Cell source | Animal | 53% |
| | Human | 28% |
| | Both | 1% |
| Culture method | 3D Encapsulation | 48% |
| | 2D Culture | 25% |
| | Both | 8% |
| No culture | N/A | 18% |
| ***In vivo* only** | | **18% (52% total)** |
| Outcome Measures | Histology | 48% |
| | Biomechanics & organization | 33% tie |
| | Gross morphology | 30% |
| | Biochemical & vascularity | 24% tie |
| | Inflammation | 20% |
| | *In Vivo* Imaging | 15% |
| Species | Rodent | 39% |
| | Other (e.g. rabbit, horse) | 11% |
| | Multiple species | 1% |
| Carrier Delivery | Supplements | 18% |
| | None | 16% |

| | | |
|---|---|---|
| | Cells | 11% |
| | Multiple carrier | 6% |
| Injury Model | Unilateral injury | 21% |
| | Bilateral injury | 18% |
| | Subcutaneous | 10% |
| | Multiple models | 2% |
| Suture | Non-resorbable (e.g. prolene) | 25% |
| | No suture | 14% |
| | Resorbable (e.g. Vicryl) | 9% |
| | Both | 3% |

**Supplemental Table 3.** Systematic review of the top clinical outcome measures from meta-analyses of randomized controlled trials of Achilles and rotator cuff tendon repair. DVT: deep vein thrombosis, ROM: range of motion, ATRS: Achilles tendon rupture score, PE: pulmonary embolism, VAS: visual analog scale.

| **Top clinical outcomes for Achilles tendon rupture** | **Percentage of n=10**[94–103] |
|---|---|
| Re-rupture & DVT | 100% |
| Wound or skin infection | 80% |
| Deep infection, return-to-sport & strength tests | 70% |
| Sural nerve injury & return-to-work | 60% |
| Tendon or skin adhesion | 50% |
| Ankle ROM & muscular (heel-rise and atrophy) | 40% |
| Tendon elongation & patient satisfaction | 30% |
| Pain, ATRS, PE, & Leppilahti score | 20% |
| Contralateral rupture, quality of life, AOFAS score, & others | 10% |
| **Top clinical outcomes for rotator cuff repair** | **Percentage of n=9**[104–112] |
| Constant, UCLA, & ASES scores | 89% |
| Re-tear | 67% |
| Pain (VAS) | 56% |
| Tendon integrity (follow-up imaging) | 44% |
| Shoulder ROM & DASH score | 33% |
| Patient satisfaction, quality of life, simple shoulder test, & frozen shoulder | 22% |
| DVT, infection, nerve injury, & others | 11% |

**Supplemental Table 4.** Guides from ISO and ASTM for testing of tissue engineered medical products (TEMPs) based on material, musculoskeletal tissue, and outcome measures.

| Testing | Reference |
|---|---|
| Biocompatibility | ISO 10993-1:2018(en), Biological evaluation of medical devices — Part 1: Evaluation and testing within a risk management process. https://www.iso.org |
| Raw Materials (TEMPs) | ASTM F2027-16, Standard Guide for Characterization and Testing of Raw or Starting Materials for Tissue-Engineered Medical Products, ASTM International, West Conshohocken, PA, 2016, www.astm.org, DOI: 10.1520/F2027-16 |
| TEMPs | ASTM F2211-13, Standard Classification for Tissue Engineered Medical Products (TEMPs), ASTM International, West Conshohocken, PA, 2013, www.astm.org, DOI: 10.1520/F2211-13 |
| Scaffolds (TEMPs) | ASTM F2150-19, Standard Guide for Characterization and Testing of Biomaterial Scaffolds Used in Regenerative Medicine and Tissue-Engineered Medical Products, ASTM International, West Conshohocken, PA, 2019, www.astm.org, DOI: 10.1520/F2150-19 |
| Hydrogels | ASTM F2900-11, Standard Guide for Characterization of Hydrogels used in Regenerative Medicine (Withdrawn 2020), ASTM International, West Conshohocken, PA, 2011, www.astm.org, DOI: 10.1520/F2900-11 |
| Collagen-Based | ASTM F3089-14, Standard Guide for Characterization and Standardization of Polymerizable Collagen-Based Products and Associated Collagen-Cell Interactions, ASTM International, West Conshohocken, PA, 2014, www.astm.org, DOI: 10.1520/F3089-14 |
| Collagen | ASTM F2212-19, Standard Guide for Characterization of Type I Collagen as Starting Material for Surgical Implants and Substrates for Tissue Engineered Medical Products (TEMPs), ASTM International, West Conshohocken, PA, 2019, www.astm.org |
| Hyaluronan | ASTM F2347-15, Standard Guide for Characterization and Testing of Hyaluronan as Starting Materials Intended for Use in Biomedical and Tissue Engineered Medical Product Applications, ASTM International, West Conshohocken, PA, 2015, www.astm.org, DOI: 10.1520/F2347-15 |
| Alginates | ASTM F2064-17, Standard Guide for Characterization and Testing of Alginates as Starting Materials Intended for Use in Biomedical and Tissue Engineered Medical Product Applications, ASTM International, West Conshohocken, PA, 2017, www.astm.org, DOI: 10.1520/F2064-17 |
| Silicone | ASTM F2042-18, Standard Guide for Silicone Elastomers, Gels, and Foams Used in Medical Applications Part II—Crosslinking and Fabrication, ASTM International, West Conshohocken, PA, 2018, www.astm.org |

| | |
|---|---|
| Absorbable (Polymers) | ASTM F2902-16e1, Standard Guide for Assessment of Absorbable Polymeric Implants, ASTM International, West Conshohocken, PA, 2016, www.astm.org, DOI: 10.1520/F2902-16E01 |
| Absorbable (Tissue Effects) | ASTM F1983-14, Standard Practice for Assessment of Selected Tissue Effects of Absorbable Biomaterials for Implant Applications, ASTM International, West Conshohocken, PA, 2014, www.astm.org, DOI: 10.1520/F1983-14 |
| Cell Viability | ASTM F2739-19, Standard Guide for Quantifying Cell Viability and Related Attributes within Biomaterial Scaffolds, ASTM International, West Conshohocken, PA, 2019, www.astm.org, DOI: 10.1520/F2739-19 |
| Cell Encapsulation (Alginate) | ASTM F2315-18, Standard Guide for Immobilization or Encapsulation of Living Cells or Tissue in Alginate Gels, ASTM International, West Conshohocken, PA, 2018, www.astm.org, DOI: 10.1520/F2315-18 |
| Biomolecules Release | ASTM F3142-16, Standard Guide for Evaluation of *in vitro* Release of Biomolecules from Biomaterials Scaffolds for TEMPs, ASTM International, West Conshohocken, PA, 2016, www.astm.org, DOI: 10.1520/F3142-16 |
| Imaging (TEMPs) | ASTM F2603-06(2012), Standard Guide for Interpreting Images of Polymeric Tissue Scaffolds, ASTM International, West Conshohocken, PA, 2012, www.astm.org, DOI: 10.1520/F2603-06R12 |
| Microstructure (TEMPs) | ASTM F2450-18, Standard Guide for Assessing Microstructure of Polymeric Scaffolds for Use in Tissue-Engineered Medical Products, ASTM International, West Conshohocken, PA, 2018, www.astm.org, DOI: 10.1520/F2450-18 |
| Wear Particles | ASTM F1904-14, Standard Practice for Testing the Biological Responses to Particles *in vivo*, ASTM International, West Conshohocken, PA, 2014, www.astm.org, DOI: 10.1520/F1904-14 |
| Tendon | ASTM F2903-11, Standard Guide for Tissue Engineered Medical Products (TEMPs) for Reinforcement of Tendon and Ligament Surgical Repair (Withdrawn 2020), ASTM International, West Conshohocken, PA, 2011, www.astm.org, DOI: 10.1520/F2903-11 |
| Cartilage | ASTM F2451-05(2010), Standard Guide for *in vivo* Assessment of Implantable Devices Intended to Repair or Regenerate Articular Cartilage (Withdrawn 2019), ASTM International, West Conshohocken, PA, 2010, www.astm.org, DOI: 10.1520/F2451-05R10 |
| Disc | ASTM F2346-18, Standard Test Methods for Static and Dynamic Characterization of Spinal Artificial Discs, ASTM International, West Conshohocken, PA, 2018, www.astm.org, DOI: 10.1520/F2346-18 |
| Disc | ASTM F2884-12, Standard Guide for Pre-clinical *in vivo* Evaluation of Spinal Fusion, ASTM International, West Conshohocken, PA, 2012, www.astm.org, DOI: 10.1520/F2884-12 |

| | |
|---|---|
| Bone | ASTM F2721-09(2014), Standard Guide for Pre-clinical *in vivo* Evaluation in Critical Size Segmental Bone Defects, ASTM International, West Conshohocken, PA, 2014, www.astm.org, DOI: 10.1520/F2721-09R14 |
| Orofacial | ASTM F1027-86(2017), Standard Practice for Assessment of Tissue and Cell Compatibility of Orofacial Prosthetic Materials and Devices, ASTM International, West Conshohocken, PA, 2017, www.astm.org, DOI: 10.1520/F1027-86R17 |
| Vascular | ASTM F3225-17, Standard Guide for Characterization and Assessment of Vascular Graft Tissue Engineered Medical Products (TEMPs), ASTM International, West Conshohocken, PA, 2017, www.astm.org, DOI: 10.1520/F3225-17 |
| Wound Closure | ASTM F2458-05(2015), Standard Test Method for Wound Closure Strength of Tissue Adhesives and Sealants, ASTM International, West Conshohocken, PA, 2015, www.astm.org, DOI: 10.1520/F2458-05R15 |
| Meniscus | ASTM F3223-17, Standard Guide for Characterization and Assessment of Tissue Engineered Medical Products (TEMPs) for Knee Meniscus Surgical Repair and/or Reconstruction, ASTM International, West Conshohocken, PA, 2017, www.astm.org, DOI: 10.1520/F3223-17 |

### Supplemental Lists

**Supplemental List 1.** Provided guides from ISO 10993 for testing biocompatibility of biomaterials.

1. Evaluation and testing within a risk management process
2. Animal welfare requirements
3. Tests for genotoxicity, carcinogenicity, and reproductive toxicity
4. Selection of tests for interactions with blood
5. Tests for *in vitro* cytotoxicity
6. Tests for local effects after implantation
7. Ethylene oxide sterilization residuals
8. Framework for identification and quantification of potential degradation products
9. Tests for irritation and skin sensitization
10. Tests for systemic toxicity
11. Sample preparation and reference materials
12. Identification and quantification of degradation products from polymeric medical devices
13. Identification and quantification of degradation products from ceramics
14. Identification and quantification of degradation products from metals and alloys
15. Toxicokinetic study design for degradation products and leachables
16. Establishment of allowable limits for leachable substances
17. Chemical characterization of materials
18. Principles and methods for immunotoxicology testing of medical devices
19. Application of risk management to medical devices

**Supplemental List 2.** Provided recommendations from ASTM F2900-11 for the use of hydrogels.

1. Biological properties
    a. Adventitious agents
        i. PCR assays and sterility tests
        ii. Bacterial endotoxin test
2. Kinetics of formation, degradation, and agent release
    a. Gelling time and rate of change of material phase
        i. Tube tile test and falling ball test
        ii. Optical turbidity and scattering
        iii. Rheology
        iv. Ultrasonic methods and dielectric spectroscopy
    b. Swelling rate (and osmotic stability, below)
        i. Equivalent solvent content study
        ii. Video microscopy
        iii. Conductivity measurement
    c. Matrix degradation
        i. Mass loss measurement
        ii. Mechanical testing
        iii. Optical and NMR spectroscopy
    d. Release rate of bioactive
        i. Diffusion chamber (for mass transport, below)
        ii. Optical spectroscopy
        iii. Biochemical analysis of aliquots
3. Physical and chemical stability
    a. Osmotic stability
    b. Mechanical properties
        i. Indentation test
        ii. Tensile/compressive testing
        iii. Rheology
        iv. Sonoelastography
    c. Cell encapsulation (and cell migration, below)
        i. Optical microscopy
        ii. Modified Boyden cell
4. Mass transport
    a. Cell migration
    b. Transport of nutrients and waste
        i. Diffusion chamber
        ii. Optical density studies
        iii. Chromatography
        iv. NMR techniques
    c. Release of bioactive

**Supplemental List 3.** Provided recommendations from ASTM F2903-11 for tendon repair[113].

1. Synthetic matrices: Polymer types, structure, degradation, and analyses
2. Natural matrices: Native matrices (decellularized), other natural matrices, cell-culture-derived, and analyses
3. Cell content: Cell performance, cell types, and analyses
4. Bioactive molecules: Source and analyses
5. Sterility and biocompatibility
6. Characterization
    a. Mechanical properties: tensile properties, suture pull-out strength, burst strength, tear strength, creep/hysteresis, fatigue durability, abrasion resistance
    b. Histology, matrix components, DNA content, metabolic activity
    c. Degradation rate
    d. Stability and shelf life
7. *In vivo* preclinical tests
    a. Biocompatibility and safety tests
    b. Animal studies for therapeutic effectiveness
        i. Time points: degradable and non-degradable matrices
        ii. Large animal models suggested
        iii. Animal model development may be necessary
        iv. Use of multiple animal models may be necessary
        v. Mechanical properties (e.g., tensile) and biologic response (e.g., histology of repair site and surrounding tissue) to determine therapeutic rationale
        vi. *In vivo* degradation (use of the repair site is preferred, but subcutaneous may provide useful information)
    c. Human cadaveric studies
        i. Surgical implantation protocols
        ii. Mechanical function at scale
8. Manufacturing
9. Clinical evaluation
    a. Prospective, randomized controlled, patient-blinded clinical trials
        i. Validated patient-reported general health-related quality of life instruments (e.g., EuroQoL, EQ-5d)
        ii. Condition/joint-specific instruments for pain (e.g. ASES score, UPenn, UCLA, or Constant shoulder scores)
        iii. Structural assessment by imaging to determine failed repair and/or gap formation
        iv. Independent assessments of joint mobility, strength level, and post-operative pain levels
10. Issues to consider
    a. Preclinical issues: Identifying appropriate design criteria, establishing useful release criteria, development of relevant animal model.
    b. Manufacturing issues: Identifying release specifications to ensure uniform product, establishing useful shelf life, convenient storage.

c. Clinical issues: Identifying optimal surgical application techniques, identification of appropriate patient populations, demonstrating effectiveness or efficacy in specific patient populations using appropriate clinical studies
d. Regulatory issues: Identifying primary function

Loose recommendation

- Degradation
- Pharmaco-kinetic and –dynamic analysis
- Study duration
- Statistics

No recommendation

- Weight
- Gender
- Follow-up arthroscopy
- Post-op MRI or radiograph
- Immune response
- Hematology
- Other toxicity tests (genotoxicity, carcinogenicity, reproductive toxicity)

### Supplemental Methods

<u>Systematic Reviews:</u> Systematic reviews were performed based on PRISMA guidelines[114]. *Search for Preclinical Outcomes of Tendon Repair with Hydrogels (preclinical articles):* We searched for peer-reviewed articles in PubMed using two search queries: 1. (tendon) AND (hydrogel) and 2. ((tendon) AND (hydrogel)) AND (scaffold) (Figure S1). *Search for Clinical Outcomes of Achilles and Rotator Cuff Repair (clinical articles):* We searched for peer-reviewed systematic reviews and meta-analyses of randomized controlled trials in PubMed using two search queries: 1. ((Achilles tendon rupture) AND (systematic review)) AND (randomized controlled trial) and 2. ((Rotator cuff repair) AND (systematic review)) AND (randomized controlled trial) (Figure S5). Results were filtered to English-language articles published from the past 10-years from 2010-2020. Searches were exported to a reference manage (Zotero) for review and duplicates were removed. Hand-selected articles were also included. *Selection criteria:* Inclusion criteria for preclinical outcomes included peer-reviewed original research of natural, synthetic, or combined natural-synthetic hydrogels using *in vitro* stem cells and/or tendon fibroblasts and/or *in vivo* tendon injury models, and exclusion criteria included non-tendon, non-hydrogel, reviews, and retracted articles. Inclusion criteria for clinical outcomes included peer-reviewed systematic reviews and meta-analyses (SMA) of randomized controlled trials (RCT) for Achilles tendon and/or rotator cuff repair, and exclusion criteria included non-tendon, non-repair, non-SMA of RCT, and retracted articles. Articles titles and abstracts were reviewed to ensure that each article met the inclusion criteria, and then full-texts were reviewed, final articles selected, and data extracted from selected articles by one reviewer (RCL). *Data extraction and analysis:* All outcome measures that were used in all selected original publications were extracted using an all-inclusive approach to avoid bias. The usage of each outcome measure from each publication was counted using a binary scoring system of 0 (not used) or 1 (used). The total usage count for each outcome measure was divided by total number of publications for each systematic review (preclinical, clinical Achilles, clinical cuff) to calculate the percent usage of each outcome measure. The percent usage was organized by the most used to least used outcome measure, and data were graphically represented using pie charts and bar graphs. No statistical comparisons were performed. *Search results:* The searches yielded 194 preclinical and 40 clinical articles (Figure S1 and S5). Twelve hand-selected articles were included in the preclinical search (Figure S1)[22,28,32,39,40,53,54,64,85–87,90]. Removal of duplicates and non-English articles yielded 154 preclinical and 39 clinical articles. Following full-text review, 87 preclinical and 19 clinical articles met the inclusion criteria (Figure S1 and S5). The top outcomes were reported in Table S2 for preclinical and Table S3 for clinical, as well as graphed in Figure S1 and S5. For preclinical articles, the usage counts of all included outcome measures were graphed in Figures S2 (hydrogels), S3 (*in vitro* outcomes), and S4 (*in vivo* outcomes). The conclusions of these systematic reviews were described in the main text.

<u>Modulus Conversion:</u> Shear storage modulus (G′) was converted to elastic Young's modulus ($E$) using rubber elasticity theory

$$E = 2G(1 + \nu),$$

where $\nu$ is the Poisson's ratio (0.5 for a hydrogel, Figure S9)[115].

*Peptide Synthesis:* Four peptides in total were synthesized. Briefly, peptides were synthesized via solid phase peptide synthesis in 0.25 mmol batches, using MBHA rink-amide resin (0.52 meq/g; NovaBiochem). Fmoc protecting groups were deprotected with 20% piperidine (Millipore Sigma, St Louis, MO) in dimethylformamide (DMF; Thermo Fisher Scientific, Waltham, MA), and amino acid residues (ChemPep, Wellington, FL) were double coupled at 5x excess at 75 °C, where each coupling reaction ran for 8 minutes and used N,N'-diisopropylcarbodiimide (DIC; 1 M in DMF; AppTech, Carlsbad, CA) and ethyl(hydroxyimino) cyanoacetate (OxymaPure; 1 M in DMF; CEM Corporation, Matthews, NC). Peptides were cleaved from the resin in a trifluoroacetic acid (TFA; Thermo Fisher Scientific, Waltham, MA)/triisopropylsilane (TIPS; Thermo Fisher Scientific, Waltham, MA)/water solution (95%/2.5%/2.5% v/v) with phenol (2.5% w/v; MilliporeSigma, Burlington, MA) for 2 hours, followed by precipitation in diethyl ether (Thermo Fisher Scientific, Waltham, MA) and 3x diethyl ether washes. Peptides were then dried overnight at room temperature prior to purification via reverse phase high performance liquid chromatography (HPLC; XBridge C18 OBD 5 µm column; Waters Corporation, Milford, MA). The two crosslinker peptides were purified using a linear gradient from 25 to 33% acetonitrile, the RGDS peptide was separated with a gradient from 22 to 28% acetonitrile, and the mfCMP was separated with a gradient from 20 to 30% acetonitrile.

*mfCMP Synthesis*: A multifunctional collagen mimetic peptide (mfCMP, $(PKG)_4PK(alloc)G(POG)_6(DOG)_4$, where O is a hydroxyproline residue) was synthesized using solid phase peptide synthesis, similar to the previously described method[116]. Here, this sequence was made in 0.1 mmol batches, using TentaGel® Resin (0.19 meq/g; Peptides International, Louisville, KY). After separation via HPLC collected fractions were frozen, lyophilized, and the identity was confirmed with mass spectrometry (Figure S6); this identified fraction was then further purified by dialysis to remove any other remaining ions, frozen, and re-lyophilized. Stock solutions were prepared in DPBS at 5 mM by weight.

*mfCMP Assembly and Characterization*: mfCMP solutions were prepared in 1x DPBS at 5 mM by weight. The solution was heated to 85 °C and held for 15 minutes to disassemble the peptide into individual peptide strands. The solution was then slowly cooled to room temperature and left to assemble for 48 hours, whereupon the assembled solutions were aliquoted, frozen, and lyophilized for use in experiments. Triple helix formation and mfCMP melting temperature were characterized according to a previously published protocol for circular dichroism (CD, Figure S8)[116].

Briefly, mfCMP solutions were prepared at 0.3 mM in DPBS, assembled as described above, and loaded into a 1 mm path length quartz cuvette. Wavelength and temperature scans were conducted on a J-1500 Circular Dichroism Spectrophotometer (JASCO Corporation, Easton, MD). Wavelength scans were taken from 250 nm to 190 nm at various temperatures (4, 25, 47, 40, and 60 °C) and averaged over 4 scans, and temperature scans were run from 4 °C to 80 °C at 225 nm wavelength. All data was converted to mean residue ellipticity [θ] (deg $cm^2$/dmol). Temperature scan data was fit to a Boltzmann curve, and the second derivative was used to determine the inflection point, and thus the melting temperature ($T_m$, where half the peptides are assembled and the other half are disassembled into single strands).

*PEG-4SH Modification:* Four-arm poly(ethylene glycol) (PEG-4OH; JenKem Technology, USA) was functionalized with thiol end groups to produce a four-arm poly(ethylene glycol) tetrathiol (PEG-4SH), according to a previously established protocol[117]. Four-arm poly(ethylene glycol) (PEG-4OH; JenKem Technology, USA) was functionalized with thiol end groups to produce a four-arm poly(ethylene glycol) tetrathiol (PEG-4SH), according to a previously established protocol[117]. Briefly, PEG-4OH ($M_n$ ~ 20,000 g/mol, 10 g) was reacted with allyl bromide (Thermo Fisher Scientific, Waltham, MA) to synthesize an allyl ether-functionalized PEG (PEG-4 Allyl Ether). Next, the allyl ether functional groups were reacted with thioacetic acid (Acros Organics, Fair Lawn, NJ) via photo-initiated radical thiol–ene click chemistry to produce PEG-4Thioacetate. Finally, the thiol group was deprotected via reaction with sodium hydroxide (NaOH; Thermo Fisher Scientific, Waltham, MA) to generate the product, PEG-4SH. Each step in the PEG modification was confirmed with $^1$H-NMR (Bruker AVIII400; Bruker Daltonics, Billerica, MA) (Figure S7). PEG-4SH was then treated with tris(2-carboxyethyl)phosphine (TCEP; Millipore Sigma, St. Louis, MO) in deionized (DI) water (~40 mL water and 350 mg TCEP per 1 g PEG-4SH) for 16 hours to reverse formation of any disulfide bonds that may have formed. The PEG-4SH was then dialyzed (MWCO 1 kDa; Spectrum Laboratories, New Brunswick, NJ) against acidified DI water (pH 4) for 24 hours, frozen, and lyophilized. 80 mM (thiol end group concentration) stock solutions of PEG-4SH in Dulbecco's phosphate-buffered saline (DPBS; Thermo Fisher Scientific, Waltham, MA) were made, where Ellman's assay was used to confirm the stock concentration.

*LAP* Synthesis: The photoinitiator used in hydrogel formation, lithium phenyl-2,4,6-trimethylbenzoylphosphinate (LAP), was synthesized according to a previously published protocol[117]. Briefly, equimolar amounts of 2,4,6-trimethylbenzoyl chloride (MilliporeSigma, Burlington, MA) and dimethylphenylphosphonite (Acros Organics, Fair Lawn, NJ) were reacted under argon for 16 hours at room temperature. Lithium bromide (4x molar excess; MilliporeSigma, Burlington, MA) in 2-butanone (MilliporeSigma, Burlington, MA) was stirred in, heated to 50 °C, and reacted for 10 minutes. The precipitate was filtered, followed by 3x rinse with 2-butanone, dried under vacuum, and the product identity was confirmed with $^1$H-NMR (Figure S7).

*Hydrogel Formulations:* The modulus of photoinitiated PEG-peptide hydrogels has been shown to be a function of the concentration of monomers in the precursor solution and the polymerization time amongst other variables: for example, with 2mM integrin binding peptide in the precursor solution, hydrogels can be formed by irradiation with low doses of long wavelength UV light (10mW/cm$^2$ at 365nm for 1 minute) for achieving a range of equilibrium swollen storage moduli from 6 to 10 wt% PEG-4SH (1:1 SH:alloc). These moduli fall within the range of those found for connective tissues (E ~ 0.5 to 5 kPa)[118]. For preliminary evaluation of hydrogels relevant for preclinical evaluation, we thus examined hydrogels on the low end of this range for cell encapsulation and 3D culture (6 wt% with and without mfCMPs) and on the high end of this range for stabilizing rotator cuff injuries (10 wt% without mfCMPs). To enable visualization within tendon defects, as detailed within the main text, here we also established a formulation of 8 wt% PEG-4SH modified with AF647 (80 µM) with and without mfCMPs, which resulted in hydrogels with *in situ* moduli similar to those of 6 wt% hydrogels without AF647.

*Equilibrium Swollen Gel Modulus Measurements:* Hydrogels (20 µl; 8 wt%; 80 µM AF647 labeling) were formed in syringe molds as described in the main text. The 0 mM mfCMP composition was prepared: 2 mM RGDS, 5.75 mM degradable crosslinker, 1.25 mM scrambled crosslinker. After gelation, hydrogels were transferred to a non-tissue culture treated 48-well plate (CELLTREAT, Pepperell, MA) with 1x DPBS (500 µL) and allowed to equilibrium swell overnight at room temperature while rocking. After conducting diameter measurements, storage modulus measurements were collected similarly to the *in situ* rheology measurements, using a Peltier plate at 25 °C in place of the quartz plate UV curing light accessory and a gap height equal to that of each hydrogel (1% strain and 2 rad/s frequency). Storage modulus was converted to Elastic modulus (Figure S9, SM; Modulus Conversion).

*rBMSC Culture*: Briefly, the epiphyses of tibiae and femurs were cut off using bone shears to expose the marrow cavities. Each diaphysis was placed within individual, sterile 0.6mL tubes. The bottom of each tube was removed using a sterile #15 scalpel blade. Each 0.6mL tube was placed concentrically inside a sterile 1.5mL tube with 0.5mL of DMEM. The concentric tubes were centrifuged at 500g for 5-minutes to pellet the bone marrow.

*alamarBlue™*: Briefly, a solution of DMEM (Phenol Red-free high glucose DMEM, VENDOR; 50 U/mL penicillin; 50 µg/mL streptomycin; 0.2% v/v Fungizone) and alamarBlue™ was prepared (1:10 alamarBlue™:DMEM). Old media was removed and cells were incubated with the alamarBlue™ solution for 4 hours at 37 °C. The solution was moved to a new 96-well plate and fluorescence measured on a plate reader.

*3D Cell Encapsulation*: Briefly, the complete growth media used for cell expansion was replaced with bFGF-free media 24 hours before encapsulation (bFGF-free media: low glucose DMEM, 50 U/mL penicillin, 50 µg/mL streptomycin, 0.2% v/v Fungizone, 10% v/v FBS). The cells were trypsinized (1x Trypsin; Thermo Fisher Scientific, Waltham, MA), added to bFGF-free media, centrifuged, and counted.

*RNA Isolation from Synthetic Hydrogels:* hMSCs were encapsulated within unlabeled hydrogels, as described in *3D Cell Encapsulation*, above. RNA was isolated and the total yield was quantified 3- and 19-days after encapsulation via a modified version of a published protocol[119] (Figure S10). Hydrogels were washed in DPBS thrice, and then individual hydrogels were placed in 2mL centrifuge tubes, frozen in liquid nitrogen, and stored at -80ºC. For RNA isolation, an autoclaved steel ball was added to each tube. Subsequently, hydrogels were pulverized for 30-seconds at 30Hz twice in liquid-nitrogen cooled blocks (MM400, Retsch, Verder Scientific). Pulverized hydrogels were incubated in Trizol-Chloroform and then pulverized again for 2-minutes at 20Hz. The final pulverized solution was transferred to Phase Lock Gel Tubes (5PRIME, Quantabio) and centrifuged at 4ºC at 14,000rpm for 5-minutes. RNA from the supernatant was isolated following the protocol of a commercially available kit (RNeasy Plus Micro Kit, Qiagen).

*Formation in Achilles Defects*: Injured Achilles tendons were placed on a chemical-resistant slippery PTFE sheet (McMaster-Carr, Elmhurst, IL) to prevent adherence of the hydrogel to the underlying surface during photopolymerization. The hydrogel solution was pipetted into the tendon defects and UV-polymerized with a collimated lamp in a step-wise fashion to limit the amount of solution that flowed through and spilled out of the defect. Specifically, either 5 or 10 µl were added to the defects (full-thickness and complete laceration, respectively) in 3 stages: first, 20% of the solution was added to the defect followed by a 2-minute irradiation; stage 1 was repeated; and finally, the remaining

60% of the hydrogel solution was pipetted into the defect and a final 4-minute irradiation was performed to create a tendon-hydrogel complex.

### Supplemental Figures

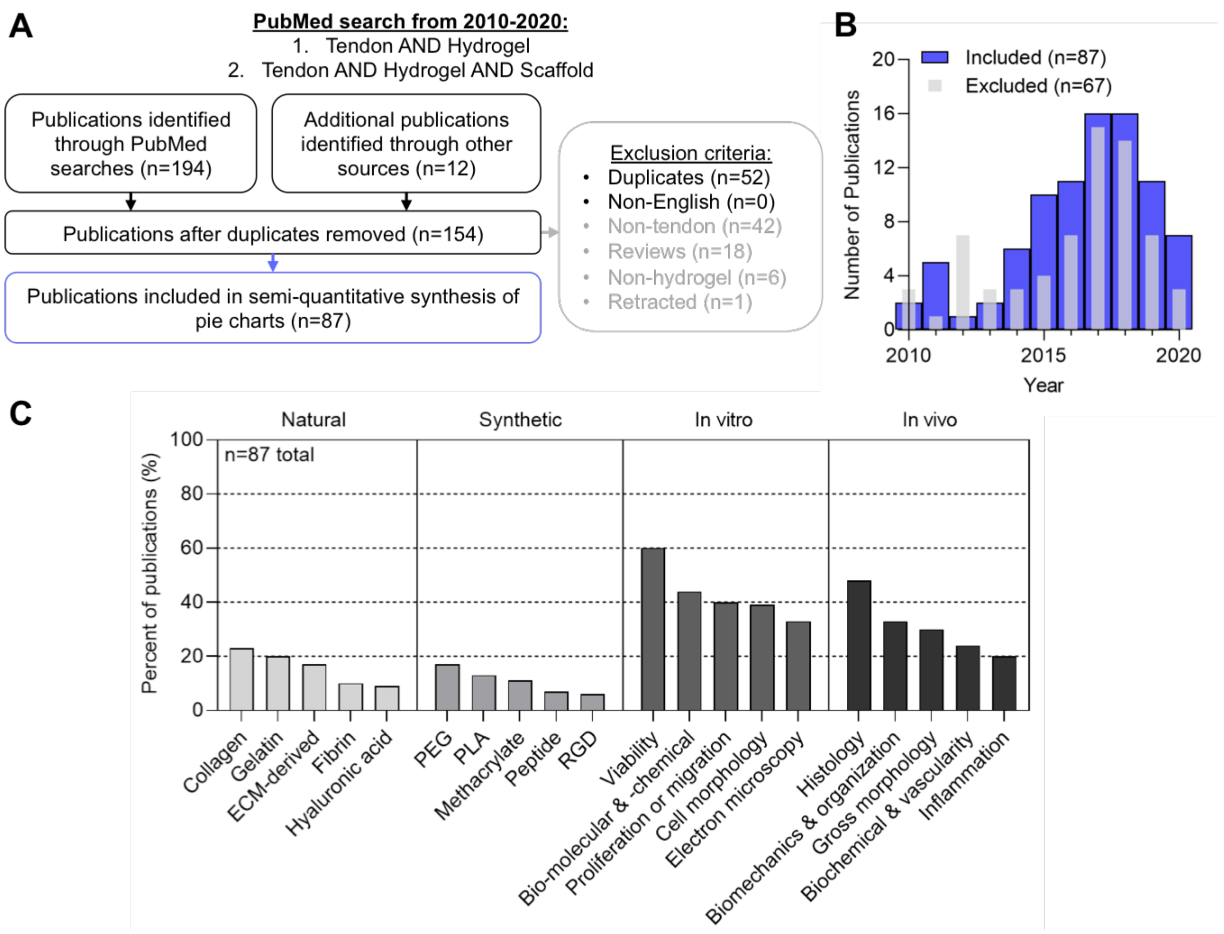

**Figure S1. Preclinical systematic review summary.** (A) PRISMA summary, (B) histogram for the number of included and excluded articles, and (C) a summary of the top 5 outcomes for natural and synthetic hydrogels, as well as for *in vitro* and *in vivo* studies.

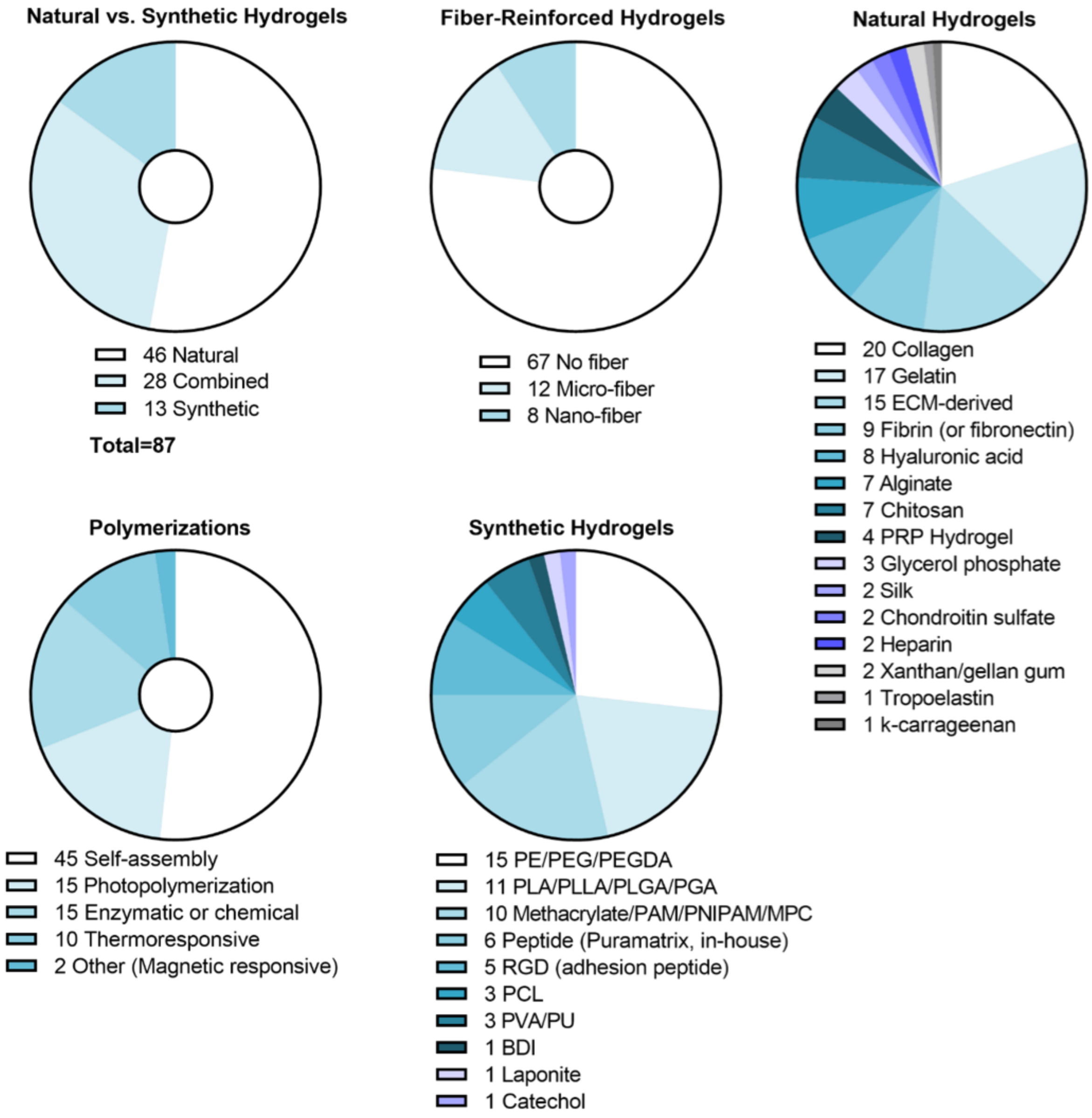

**Figure S2. Preclinical systematic review: hydrogel types.** Data presented as pie charts of usage counts for each outcome.

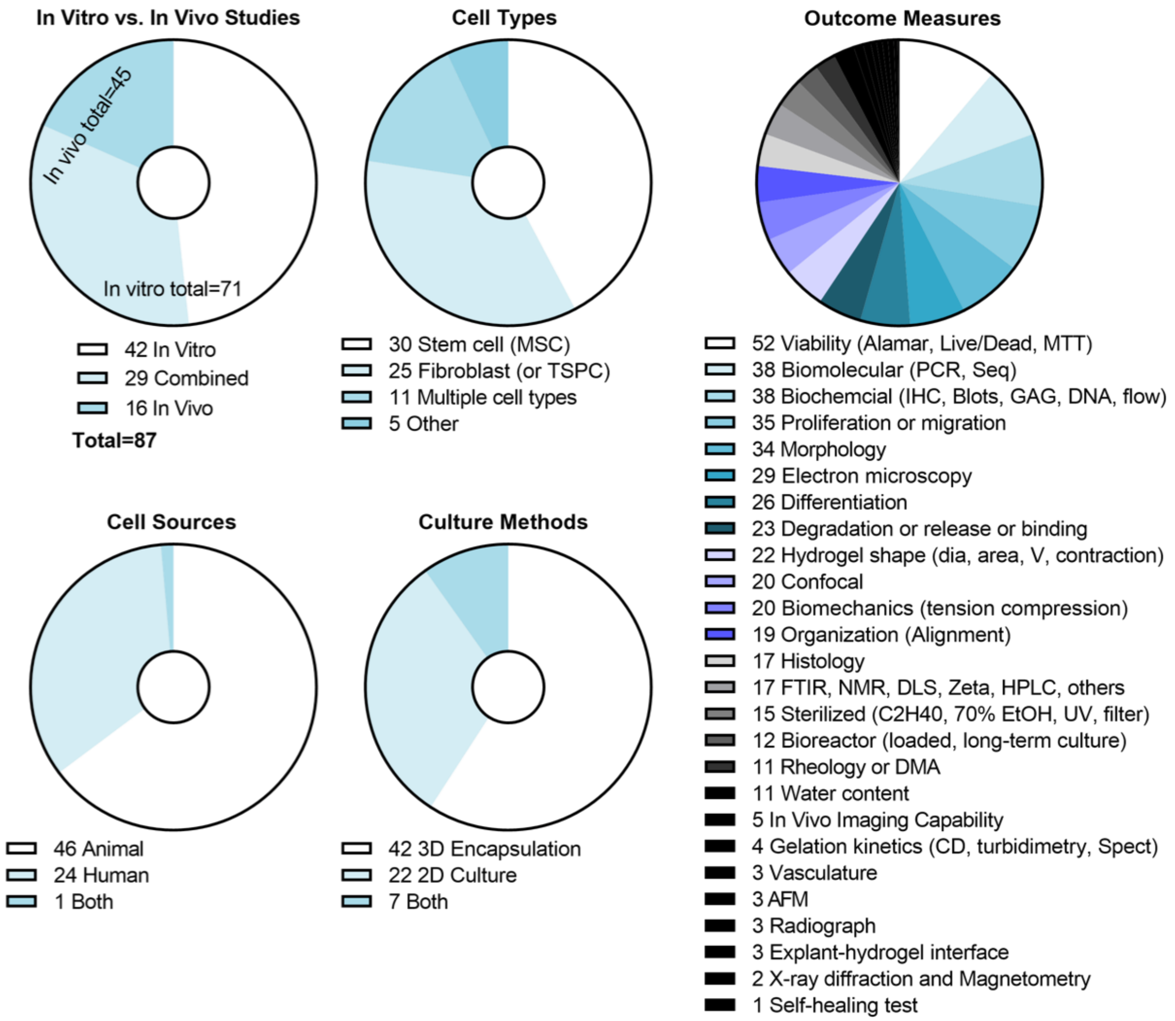

**Figure S3. Preclinical systematic review: *In vitro* outcomes.** Data presented as pie charts of usage counts for each outcome.

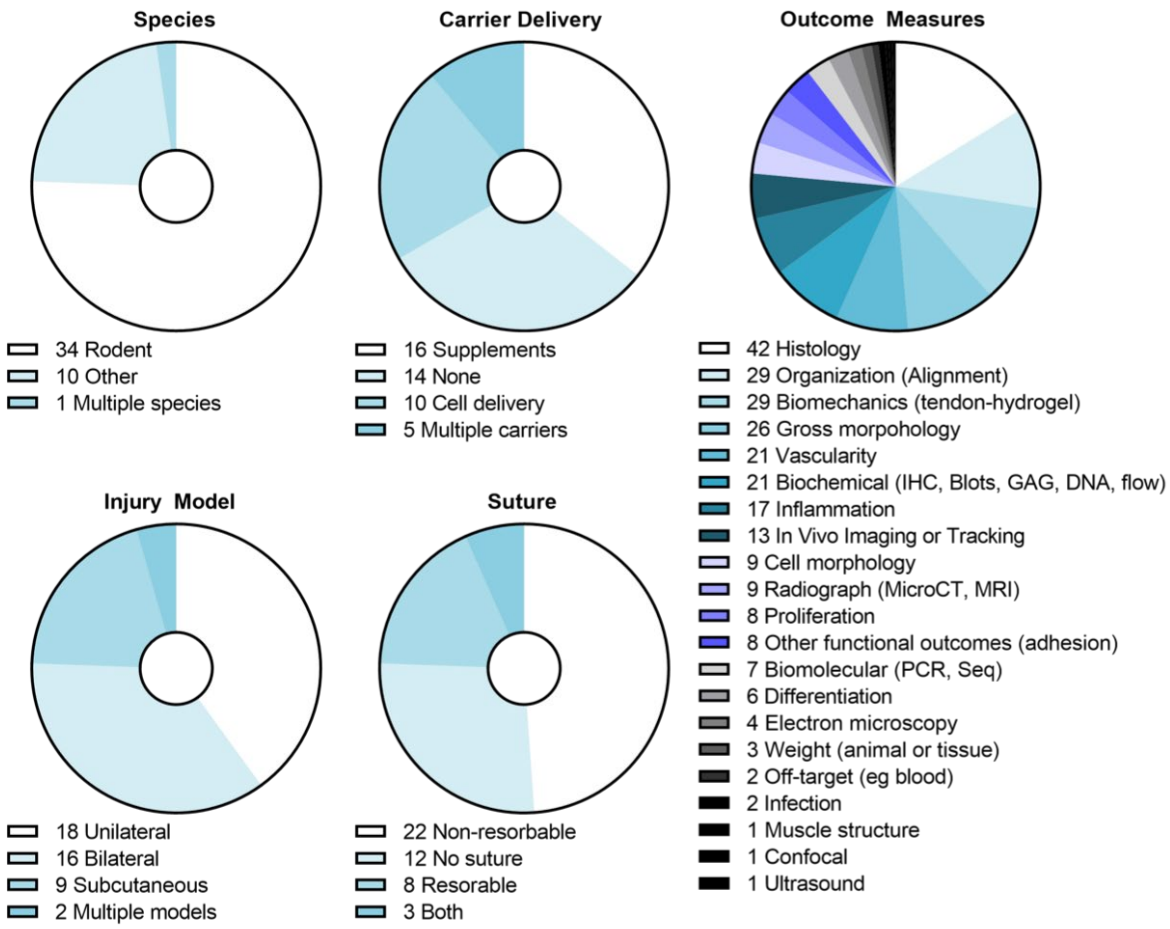

**Figure S4. Preclinical systematic review: *In vivo* outcomes.** Data presented as pie charts of usage counts for each outcome.

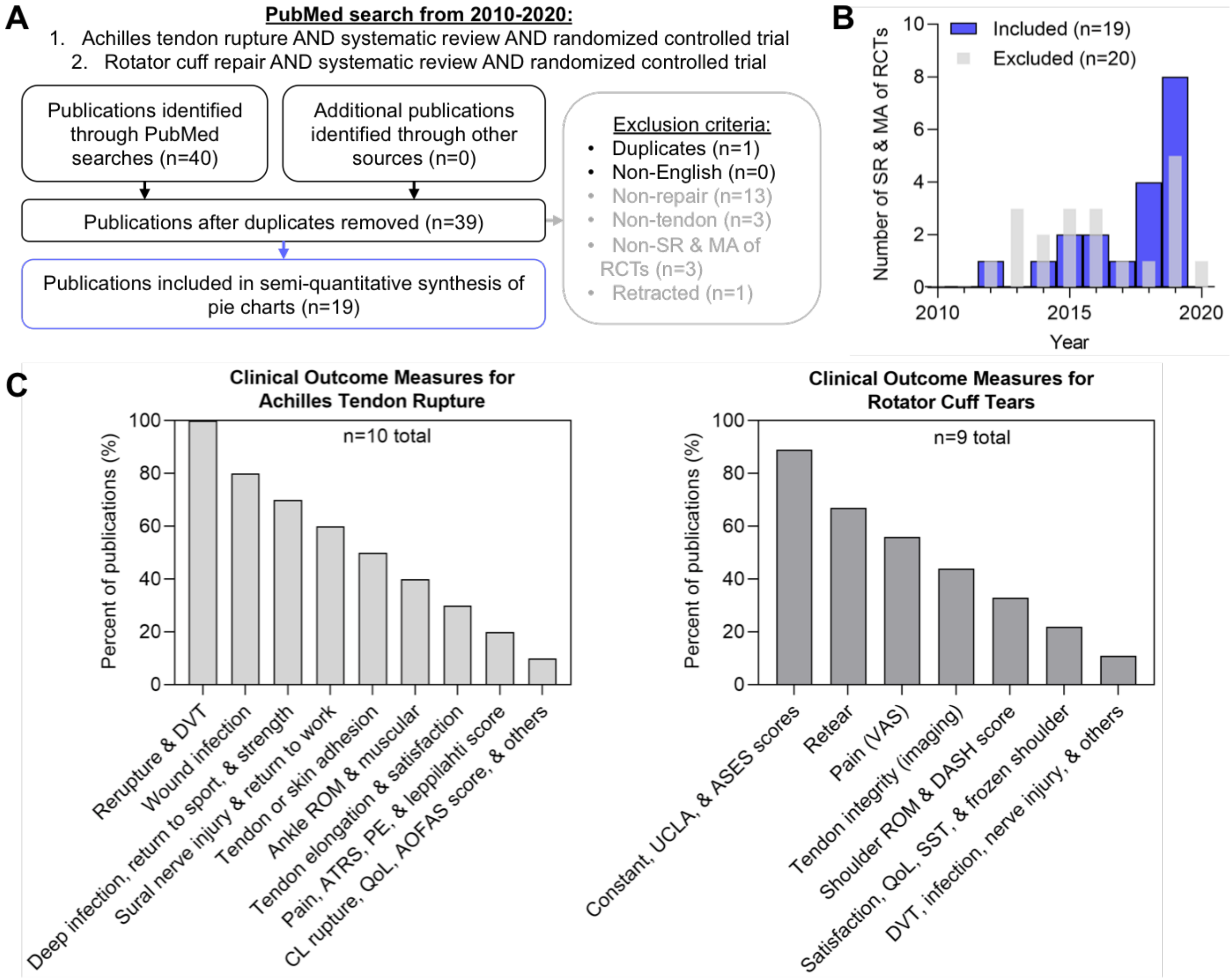

**Figure S5. Clinical systematic review summary.** (A) PRISMA summary, (B) histogram for the number of included and excluded articles, and (C) a summary of the top clinical outcome measures used for Achilles tendon rupture and rotator cuff tears.

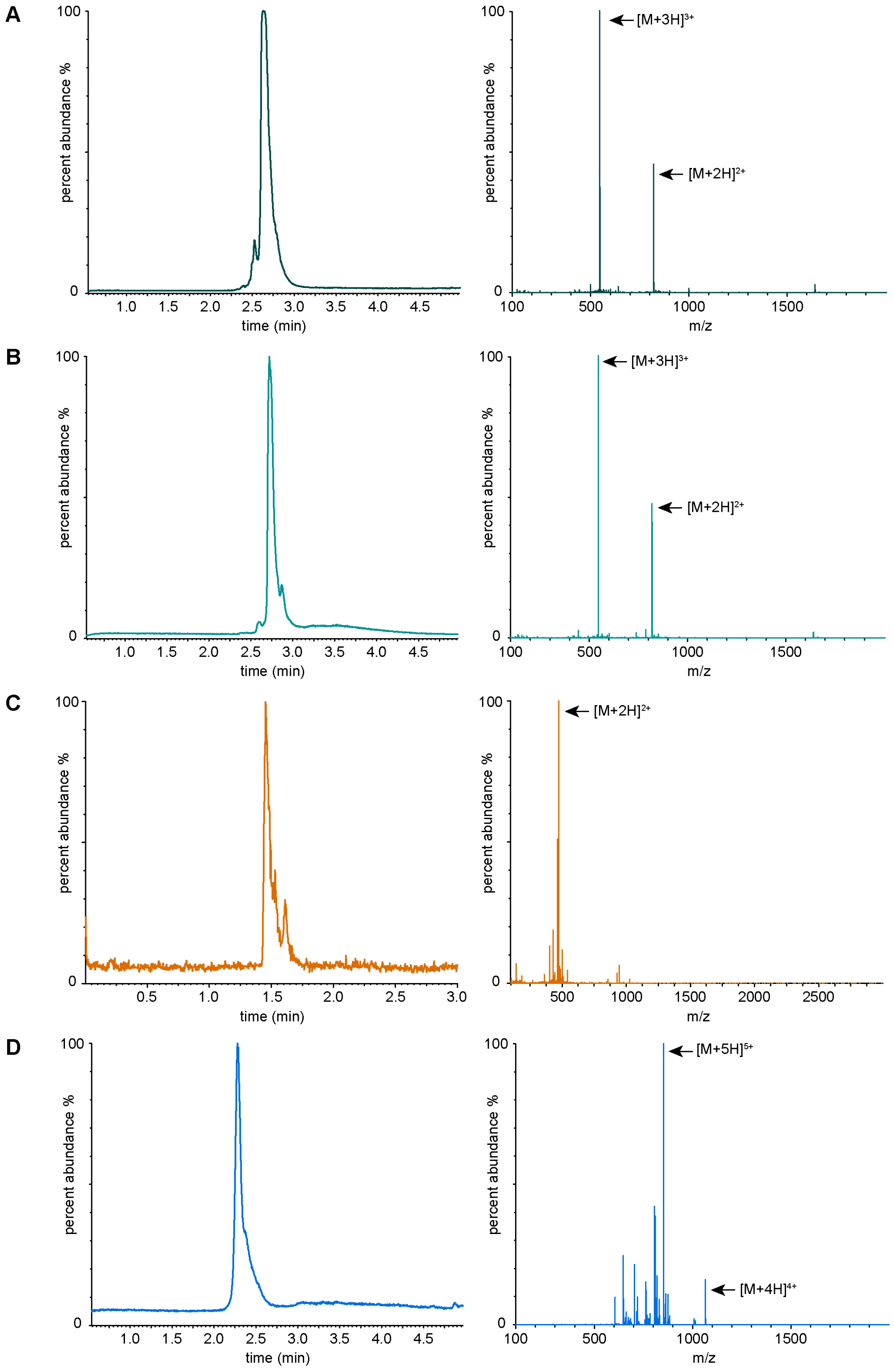

**Figure S6. Mass spectrometry results.** (A) Xevo G2-S QTof (XEVO) UPLC trace and electrospray ionization (ESI+) mass spectrometry of scrambled linker peptide (KK(alloc)GIQPGGWGQGK(alloc)K). The desired product was observed with the expected molecular weight of 1635 g/mol ($[M+2H]^{2+}$ = 818 g/mol and $[M+3H]^{3+}$ = 546 g/mol). (B) XEVO UPLC trace and electrospray ionization (ESI+) mass spectrometry of degradable linker peptide (KK(alloc)GGPQG↓IWGQGK(alloc)K). The desired product was observed with the expected molecular weight of 1635 g/mol ($[M+2H]^{2+}$ = 818 g/mol and $[M+3H]^{3+}$ = 546 g/mol). (C) Single Quadruple Detector 2 (SQD2) UPLC trace and electrospray ionization (ESI+) mass spectrometry of integrin-binding peptide RGDS (K(alloc)GWGRGDS). The desired product was observed with the expected molecular weight of 945 g/mol ($[M+2H]^{2+}$ = 473 g/mol). (D) XEVO UPLC trace and electrospray ionization (ESI+) mass spectrometry of mfCMP ($(PKG)_4PK(alloc)G(POG)_6(DOG)_4$). The desired product was observed with the expected molecular weight of 4256 g/mol ($[M+4H]^{4+}$ = 1065 g/mol and $[M+5H]^{5+}$ = 852 g/mol).

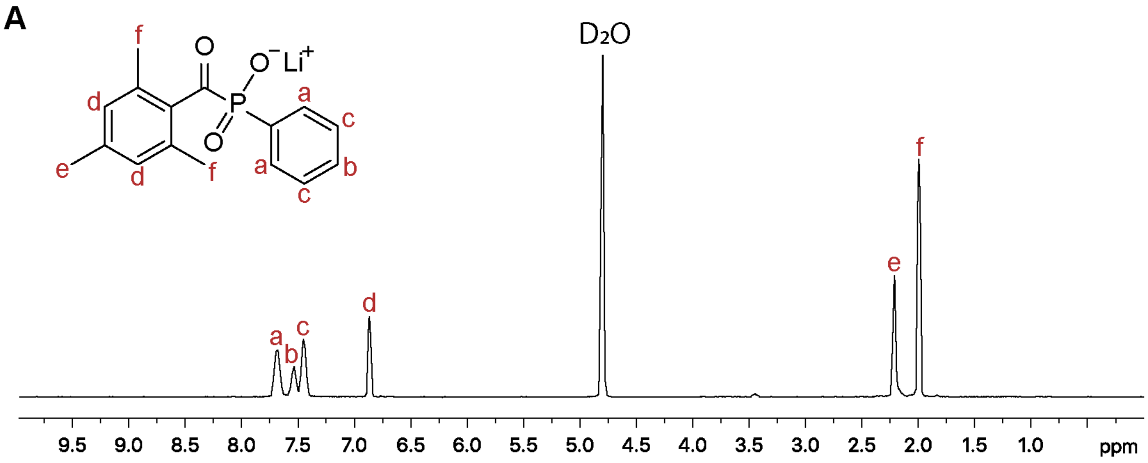

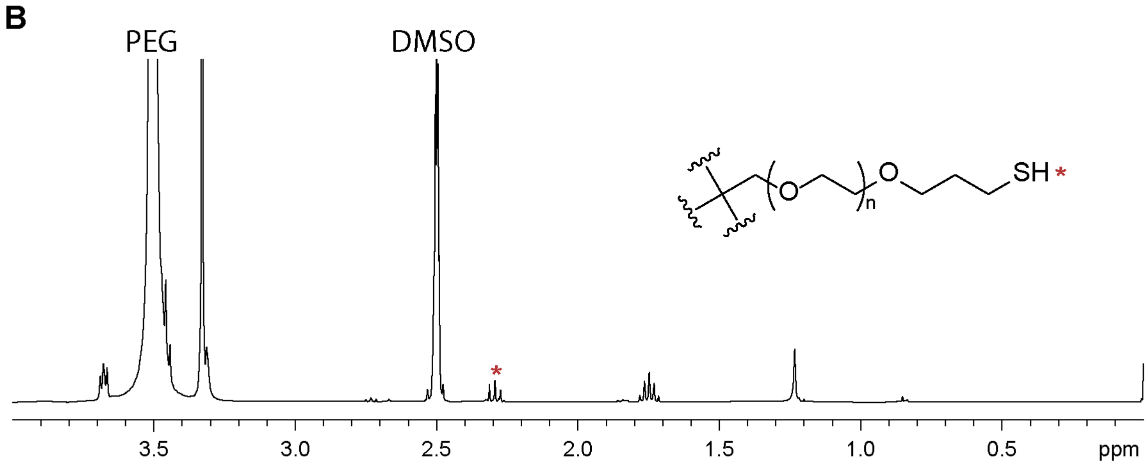

**Figure S7. $^1$H NMR results.** (A) $^1$H NMR spectrum of LAP photoinitiator in $D_2O$ (400 MHz). (B) $^1$H NMR spectrum of PEG-4SH in DMSO-d6 (400 MHz). The peak representing the thiol proton (starred) was integrated and normalized to the integration of the peak representing the PEG backbone protons to determine thiol functionality in product (~98%).

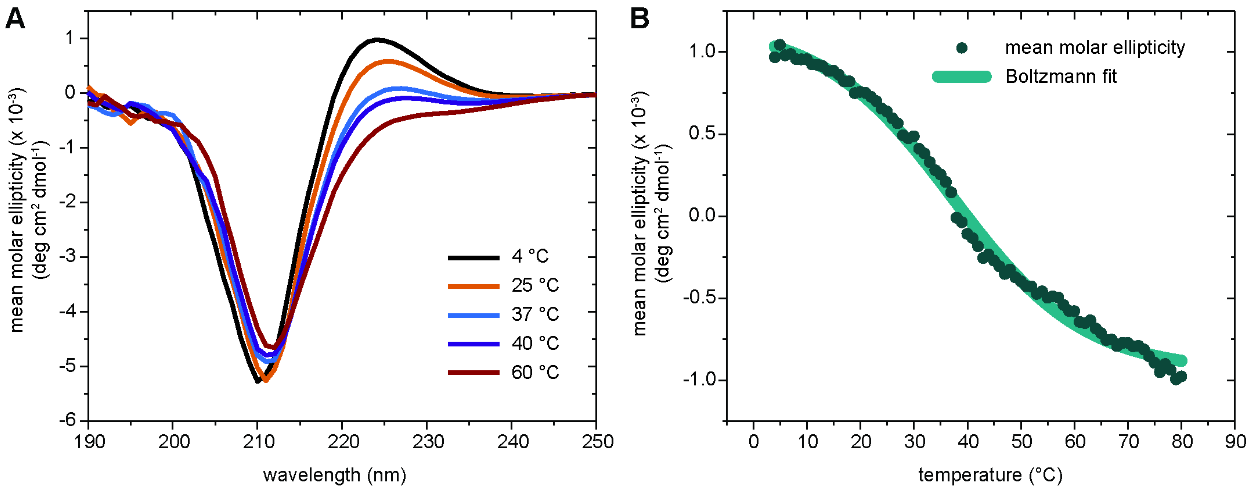

**Figure S8. Circular dichroism.** (A) Wavelength scans of mfCMP at increasing temperatures (0.3 mM mfCMP in 1x DPBS). The decrease in the 225 nm peak demonstrates melting of the triple helix. (B) Temperature scans of mfCMP at 225 nm (0.3 mM mfCMP in 1x DPBS). Melting temperature ($T_m$, point at which 50% of the peptide is no longer in the triple helical conformation) was determined to be ~37 °C.

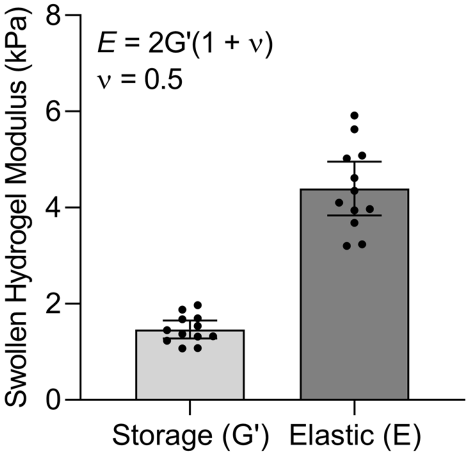

**Figure S9. Equilibrium swollen modulus.** Storage modulus was measured on hydrogels (here, without mfCMP) that had been allowed to equilibrium swell overnight. Storage modulus was then converted to Elastic modulus, which falls within the range of developing tendon.

RNA Isolation from Synthetic Hydrogels

- Encapsulate and culture cells within hydrogels for culture period
- Rinse hydrogels 3x in sterile 1x DPBS
- Place individual gels in 2mL tubes
- Flash freeze in liquid nitrogen and store at -80ºC

Pulverization of Hydrogels

- Add one, autoclaved steel bead to each tube
- Place back into $LN_2$
- Using $LN_2$-cooled blocks, pulverize gels with beads 2x at 30Hz for 30sec
- Isolate RNA using Trizol-Chloroform
- Pulverize again 2x at 20Hz for 2min
- Separate using phase-lock gel tubes
- Follow micro-prep kit to yield RNA

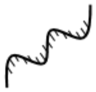

| Day | Group | [RNA] ng/µL | Total RNA (ng) |
|---|---|---|---|
| 3 | 0mM CMP | 45.5 | 544.8 |
| 19 | 0mM CMP | 36.8 | 441.6 |
| 3 | 2.5mM CMP | 33 | 396 |
| 19 | 2.5mMCMP | 30.9 | 370.8 |

**Figure S10.** Summarized RNA isolation protocol and individual hydrogel yields using this protocol.

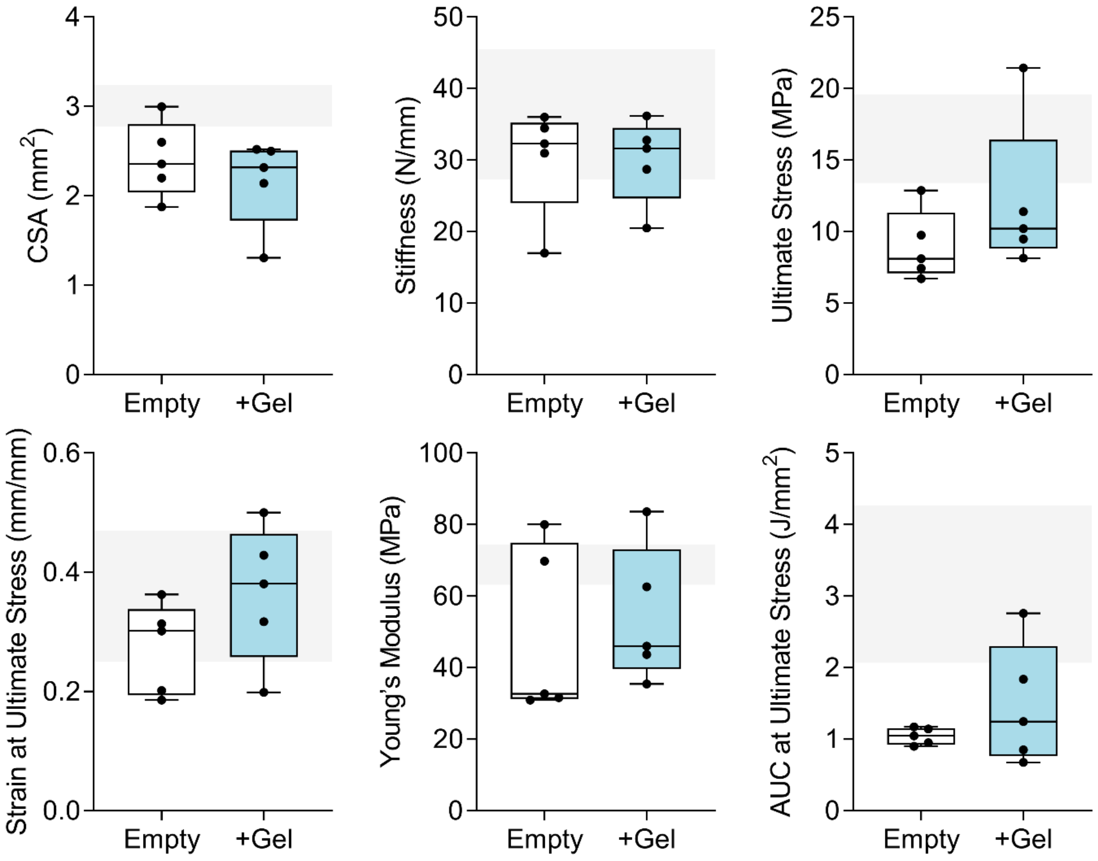

**Figure S11. Additional tensile mechanical properties.** Cross-sectional area (CSA), stiffness, ultimate stress, strain at ultimate stress, Young's modulus, and area under the curve (AUC) at ultimate stress of empty and hydrogel-filled defects with respect to non-injured tendon (intact healthy attachment) and paired-limb comparisons were made between empty and hydrogel groups. Data are presented as median ± 95% confidence intervals. Black bars: significant difference between groups ($p<0.05$).

24. Chiou, G.J., Crowe, C., McGoldrick, R., Hui, K., Pham, H., and Chang, J. Optimization of an injectable tendon hydrogel: the effects of platelet-rich plasma

and adipose-derived stem cells on tendon healing in vivo. Tissue Eng Part A. United States, **21**, 1579, 2015.

30. Tokunaga, T., Ide, J., Arimura, H., Nakamura, T., Uehara, Y., Sakamoto, H., et al. Local Application of Gelatin Hydrogel Sheets Impregnated With Platelet-Derived

Growth Factor BB Promotes Tendon-to-Bone Healing After Rotator Cuff Repair in Rats. Arthrosc J Arthrosc Relat Surg Off Publ Arthrosc Assoc N Am Int Arthrosc Assoc. United States, **31**, 1482, 2015.

98. El-Akkawi, A.I., Joanroy, R., Barfod, K.W., Kallemose, T., Kristensen, S.S., and Viberg, B. Effect of Early Versus Late Weightbearing in Conservatively Treated

Acute Achilles Tendon Rupture: A Meta-Analysis. J Foot Ankle Surg. **57**, 346, 2018.

104. Millett, P.J., Warth, R.J., Dornan, G.J., Lee, J.T., and Spiegl, U.J. Clinical and structural outcomes after arthroscopic single-row versus double-row rotator cuff

repair: a systematic review and meta-analysis of level I randomized clinical trials. J Shoulder Elbow Surg. **23**, 586, 2014.
